# Polymersomes decorated with SARS-CoV-2 spike protein receptor binding domain elicit robust humoral and cellular immunity

**DOI:** 10.1101/2021.04.08.438884

**Authors:** Lisa R. Volpatti, Rachel P. Wallace, Shijie Cao, Michal M. Raczy, Ruyi Wang, Laura T. Gray, Aaron T. Alpar, Priscilla S. Briquez, Nikolaos Mitrousis, Tiffany M. Marchell, Maria Stella Sasso, Mindy Nguyen, Aslan Mansurov, Erica Budina, Ani Solanki, Elyse A. Watkins, Mathew R. Schnorenberg, Andrew C. Tremain, Joseph W. Reda, Vlad Nicolaescu, Kevin Furlong, Steve Dvorkin, Shann S. Yu, Balaji Manicassamy, James L. LaBelle, Matthew V. Tirrell, Glenn Randall, Marcin Kwissa, Melody A. Swartz, Jeffrey A. Hubbell

**Affiliations:** Pritzker School of Molecular Engineering, University of Chicago, Chicago, IL 60637, United States; Committee on Immunology, University of Chicago, Chicago, IL 60637, United States; Animal Resources Center, University of Chicago, Chicago, IL 60637, United States; Department of Microbiology, Howard T. Ricketts Laboratory, University of Chicago, Chicago, IL 60637, United States; Department of Microbiology and Immunology, University of Iowa, Iowa City, IA 52242, United States; Department of Pediatrics, University of Chicago Comer Children’s Hospital, Chicago, IL 60637, United States; Materials Science Division, Argonne National Laboratory, Lemont, IL 60439, United States; Ben May Department of Cancer Research, University of Chicago, Chicago, IL 60637, United States; Committee on Cancer Biology, University of Chicago, Chicago, IL 60637, United States

## Abstract

A diverse portfolio of SARS-CoV-2 vaccine candidates is needed to combat the evolving COVID-19 pandemic. Here, we developed a subunit nanovaccine by conjugating SARS-CoV-2 Spike protein receptor binding domain (RBD) to the surface of oxidation-sensitive polymersomes. We evaluated the humoral and cellular responses of mice immunized with these surface-decorated polymersomes (RBD_surf_) compared to RBD-encapsulated polymersomes (RBD_encap_) and unformulated RBD (RBD_free_), using monophosphoryl lipid A-encapsulated polymersomes (MPLA PS) as an adjuvant. While all three groups produced high titers of RBD-specific IgG, only RBD_surf_ elicited a neutralizing antibody response to SARS-CoV-2 comparable to that of human convalescent plasma. Moreover, RBD_surf_ was the only group to significantly increase the proportion of RBD-specific germinal center B cells in the vaccination-site draining lymph nodes. Both RBD_surf_ and RBD_encap_ drove similarly robust CD4^+^ and CD8^+^ T cell responses that produced multiple Th1-type cytokines. We conclude that multivalent surface display of Spike RBD on polymersomes promotes a potent neutralizing antibody response to SARS-CoV-2, while both antigen formulations promote robust T cell immunity.

## INTRODUCTION

COVID-19, the disease caused by the novel coronavirus SARS-CoV-2, emerged in late 2019 and was declared a pandemic by the World Health Organization in March 2020. Since its emergence, researchers across the world have sought to rapidly develop vaccine candidates, some of which have received Emergency Use Authorization by the U.S. Food and Drug Administration^1,2^. While the first vaccines that entered the clinic were based on nucleic acid technologies, subunit vaccines are gaining attention and have also shown promise in clinical trials^3,4^. The primary antigens used in preclinical and clinical vaccine candidates are the Spike protein and its constituent receptor-binding domain (RBD). The RBD of the Spike protein binds to the ACE-2 receptor on host cell surfaces, enabling viral entry into the host cell^5,6^.

Several highly potent neutralizing antibodies have been isolated that target RBD and prevent viral binding and uptake, making it an attractive vaccine target^7–10^. Since RBD is smaller (∼25 kDa) and more stable than the full homotrimeric Spike fusion protein (∼180 kDa), it is also advantageous from a manufacturing and distribution perspective^11^. However, RBD has been shown to have lower immunogenicity than the full Spike protein or its RBD-containing S1 domain^12,13^. Materials science and engineering approaches, particularly strategies involving nanotechnology, may improve RBD immunogenicity and thus aid in the development of next-generation vaccines^14–16^. Indeed, several approaches of self-assembling RBD into virus-like particles have resulted in potent neutralizing antibody responses^17–20^.

In order to offer robust protection from infection, cellular in addition to humoral responses are needed^21–23^. Almost all convalescent individuals show T cell immunity, and the majority have both CD4^+^ and CD8^+^ SARS-CoV-2-specific T cells^24–27^. Conversely, severe disease is associated with lymphopenia and reduced T cell function^28–30^. Furthermore, T cell immunity may be more durable than humoral responses, and T cells are expected to play an important role in immune memory^23,28,31^. Therefore, the goal of this study was to improve both humoral and cellular immunogenicity of RBD and compare the efficacy of engineered nanoparticle formulations in order to inform the design of next-generation nanovaccines.

We have previously reported the development of polymersomes (PS) that self-assemble from the oxidation-responsive block copolymer poly(ethylene glycol)-bl-poly(propylene sulfide) (PEG-PPS)^32^ and shown their efficacy in delivering antigen and adjuvant to dendritic cell endosomes^33^. In endolysosomal compartments, the PPS block becomes oxidized, which initiates the restructuring of the PS into micelles and concurrent release of encapsulated payload^33,34^. These vaccine nanocarriers have been shown to activate dendritic cells, induce robust T cell immunity, and elicit high antibody titers with broad epitope coverage^33,35,36^.

In this study, we hypothesized that we could further improve the humoral responses elicited by PS while retaining their ability to induce T cell immunity by engineering them to mimic the physical form of a viral particle through multivalent surface display of antigen. We envisaged that multivalent surface display of RBD would result in enhanced crosslinking and clustering of B cell receptors (BCRs) and subsequent production of neutralizing antibodies. Here, we report on the development and preclinical evaluation of PS displaying surface-bound RBD (RBD_surf_) and PS encapsulating RBD (RBD_encap_) adjuvanted with monophosphoryl lipid A-encapsulated PS (MPLA PS). We show that mice vaccinated with RBD_surf_ in combination with MPLA PS in a prime-boost schedule develop high titers of SARS-CoV-2-neutralizing antibodies with robust germinal center responses as well as CD4^+^ and CD8^+^ T cell immunity, thus meeting our design criteria.

## RESULTS

### Formulated polymersomes exhibit long-term stability and *in vitro* activity

Having previously encapsulated antigen into PS as nanovaccines^33^, here we developed a conjugation strategy to attach antigens to their surface. To create a modular platform that could be generalized to any antigen, we synthesized N_3_-PEG-PPS (**Suppl. Fig. S1**), which, when formulated into PS, yields particles displaying clickable surface moieties (**Fig. 1a**). Upon the addition of RBD conjugated to a DBCO-containing linker, we generated PS decorated with RBD (RBD_surf_, **Suppl. Fig. S2**). We also synthesized PEG-PPS (**Suppl. Fig. S3**) and formulated PS encapsulating RBD (RBD_encap_) or adjuvant (MPLA PS, **Fig. 1a**). The loading capacities of RBD_surf_ and RBD_encap_ were 1.57% and 1.75%, respectively, comparable to previous reports of encapsulated ovalbumin^33,35^, while the loading capacity of MPLA PS was 6.46% (**Suppl. Table S1**). We confirmed the vesicular structure of PS through cryo-electron microscopy (cryoEM) and demonstrated that the different formulations have similar sizes and morphologies (**Fig. 1b, Suppl. Fig. S4**). According to dynamic light scattering (DLS) measurements, the average PS diameter is around 150 nm (**Fig. 1c,d**), which is similar to the reported size of SARS-CoV-2 particles (60-140 nm)^37^.The polydispersity index (PDI) of each formulation was < 0.2, indicative of a relatively homogenous population of nanoparticles. As indicated by their consistent size and PDI, in addition to the absence of free RBD released into solution, PS remain stable at 4 °C for at least 180 days, which can be beneficial for distribution and shelf-life considerations (**Fig. 1c, Suppl. Fig. S5**).

**Figure 1.**
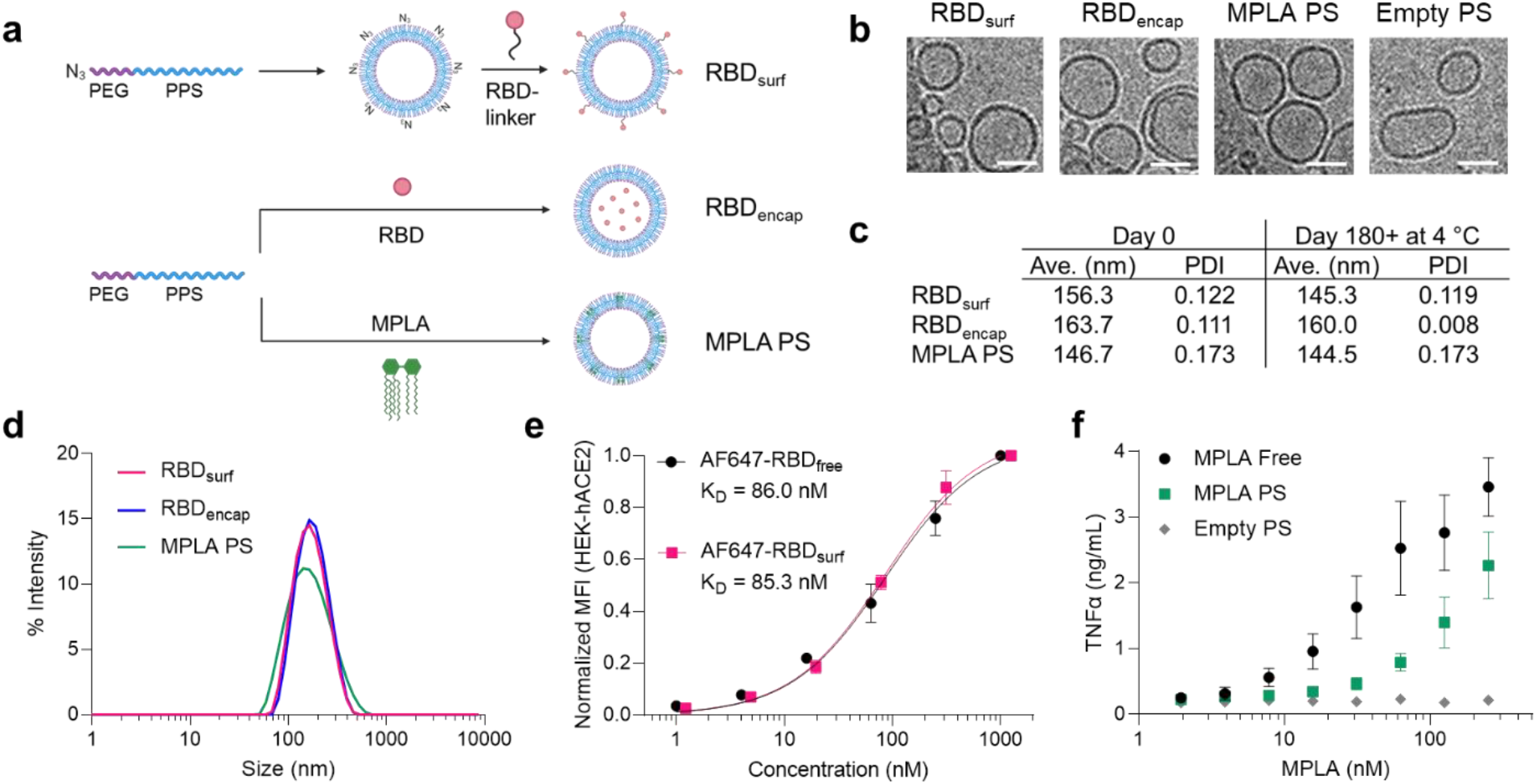
RBD and MPLA are formulated into stable, biologically active polymersomes (PS). **a**, Schematic of formulation of PS. RBD was conjugated to the surface (RBD_surf_) or encapsulated inside (RBD_encap_) of PS, and MPLA was encapsulated in the vesicle membrane (MPLA PS) due to its hydrophobicity. **b**, Representative cryo-electron microscopy images of PS, depicting vesicle structure. Scale = 50 nm. **c**, Size and polydispersity index (PDI) from dynamic light scattering (DLS) measurements of PS upon formulation and after > 6 months at 4 °C. **d**, Representative DLS curves of PS. **e**, Normalized mean fluorescence intensity (MFI) of AF647 conjugated to free RBD or RBD_surf_ by flow cytometry showing concentration-dependent binding to HEK-293 cells that express human ACE-2 (HEK-hACE2). Nonlinear regression was used to model data assuming specific binding to one site to determine equilibrium dissociation constants. **f**, Dose-dependent secretion of TNFα from cultured murine bone marrow-derived dendritic cells (BMDCs) stimulated by free MPLA, MPLA PS, or empty PS. Data represent mean ± SD for n = 2 (e) or 3 (f) replicates.

We next characterized the biological activity of the PS formulations *in vitro*. To confirm that RBD structure is not substantially altered upon conjugation to the PS surface, we quantified its ability to bind to HEK-293 cells that express human ACE-2 (HEK-hACE2, **Fig. 1e**). The normalized mean fluorescence intensity (MFI) versus RBD concentration curves were used to calculate the equilibrium dissociation constants (K_D_) for free RBD and RBD_surf_ conjugated to AF647 (AF647-RBD_free_ and AF647-RBD_surf_, respectively). The curves and K_D_ values are in excellent agreement, indicating that surface conjugation to PS did not impact ACE-2 binding of RBD. Empty PS conjugated to AF647 did not bind to HEK-hACE2, and neither PS formulation bound to HEK-293 cells lacking hACE-2 (**Suppl. Fig. S6**). Next, we confirmed that MPLA retained its ability to serve as a TLR4 agonist upon formulation in PS with a HEK-Blue™ TLR4 reporter cell line (**Suppl. Fig. S7**). To further validate MPLA PS activity in a more physiologically-relevant model, we stimulated murine bone marrow-derived dendritic cells (BMDCs) with free MPLA, MPLA PS, or empty PS, and we measured the subsequent secretion of the pro-inflammatory cytokines TNFα, IL-6, IL-1α, and IL-1β (**Fig. 1f, Suppl. Fig. S8**). For each cytokine, there was a dose-dependent increase in secretion for free MPLA and MPLA PS with only background levels of secretion for empty PS, indicating that MPLA PS successfully activated antigen presenting cells (APCs) *in vitro*. Thus, we successfully synthesized two RBD formulations of PS in addition to MPLA PS and showed that they are homogenous vesicular structures with long-term stability and *in vitro* biological activity.

### All adjuvanted formulations elicit RBD-specific IgG responses

Having confirmed that antigen- and adjuvant-loaded PS exhibit their expected bioactivity *in vitro*, we next evaluated their ability to enhance humoral and cellular immunity in mice compared to RBD_free_. We immunized mice via s.c. injection in the hocks in a prime-boost schedule 3 weeks apart and monitored antibody titers weekly (**Fig. 2a**). The total RBD-specific IgG is represented by the area under the log-transformed ELISA absorbance curves (AUC), starting at a plasma dilution of 10^−2^ (see Methods, **Suppl. Fig. S9**). All adjuvanted groups had significant anti-RBD binding antibody responses within a week after their first dose, with RBD_encap_ stimulating the highest responses (**Fig. 2b**). The antibody responses in adjuvanted groups either increased gradually or remained constant until a week after the booster, when the mean AUC increased 1.3- to 1.6-fold.

**Figure 2.**
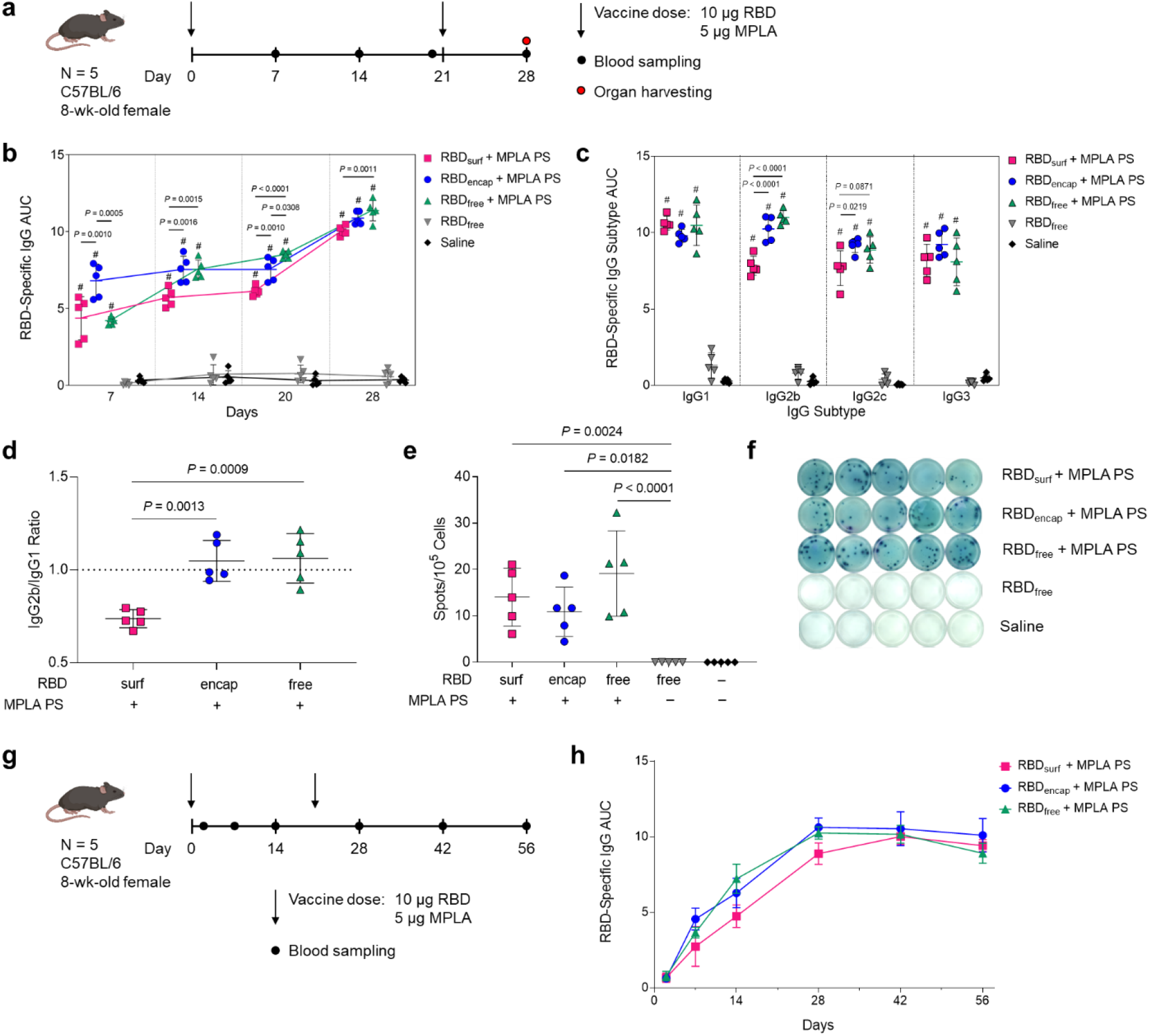
High levels of RBD-specific IgG antibodies are produced upon PS vaccination. **a**, Vaccination schedule consisting of a priming dose followed by a booster 3 weeks later. **b**, Total RBD-specific IgG antibodies over time reported as the area under the log-transformed curve (AUC) of absorbance vs. dilution. **c**, Comparison of RBD-specific IgG isotypes (IgG1, IgG2b, IgG2c, IgG3) on day 28. **d**, Ratio of AUCs of IgG2b:IgG1 isotypes. **e**, Quantification of RBD-specific IgG+ antibody secreting cells by ELISpot of splenocytes (Dunn’s post-test compared to unadjuvanted RBD_free_). **f**, Representative ELISpot wells from (e). Data plotted as mean ± SD and represent 1 of 2 experiments with n = 5 mice each. Symbols represent individual mice. **g**, Vaccine and blood sampling schedule of long-term kinetics study. **h**, Total RBD-specific IgG antibodies over time for the vaccination schedule in (g). Data represent mean ± SD for n = 5 mice. Comparisons were made using one-way ANOVA with Tukey’s post-test unless stated otherwise. ^#^ *P* < 0.0001 compared to unadjuvanted RBD_free_.

In order to explore the humoral response in further detail, we then evaluated IgG subtypes of induced antibodies at the study endpoint (d28, 1 week post-boost). While plasma antibody levels of all adjuvanted groups were similar for IgG1 and IgG3, RBD_surf_ elicited significantly lower IgG2b and IgG2c antibody responses (**Fig. 2c**). The ratio of IgG2b/IgG1 was then taken as an indication of a Th1/Th2-mediated response^38^. While RBD_encap_ and RBD_free_ + MPLA PS have a ratio of around 1, indicating a balanced Th1/Th2 response, RBD_surf_ shows a lower ratio of IgG2b to IgG1 indicating a slightly Th2-skewed response (**Fig. 2d**). Since IgA is important for combating respiratory viruses at the mucosal sites, we also measured plasma levels of RBD-specific IgA antibodies^39,40^. Although we did not expect high titers of circulating IgA from a s.c. administered vaccine, there were detectable anti-RBD IgA antibodies in all adjuvanted groups, which were significantly higher than the background-level responses elicited by unadjuvanted RBD_free_ (**Suppl. Fig. S10**).

Next, to determine if the higher antibody responses of adjuvanted groups stemmed from an expanded number of RBD-specific antibody secreting cells (ASCs), we performed an *ex vivo* RBD enzyme-linked immunosorbent spot (ELISpot) assay with splenocytes harvested 1 week post-boost. All groups receiving adjuvanted RBD showed significantly higher RBD-specific IgG^+^ ASCs compared to unadjuvanted RBD_free_, consistent with plasma antibody levels (**Fig. 2e,f**).

Finally, we evaluated the kinetics and durability of the humoral response to demonstrate the persistence of elicited antibodies (**Fig. 2g**). The RBD-specific IgG AUC for all adjuvanted groups increased until 1 week post-boost and then remained constant over the subsequent 4 weeks, indicating that the antibody responses stimulated by these vaccine formulations persist for at least 2 months in mice after the initial dose. Taken together, MPLA PS-adjuvanted RBD_surf_, RBD_encap_, and RBD_free_ all stimulated persistent anti-RBD antibodies and increased the frequencies of RBD-specific ASCs in the spleen.

### RBD-surface-decorated polymersomes, but not RBD-encapsulated polymersomes, induce neutralizing antibodies

After analyzing the quantity of RBD-specific antibodies produced by the vaccine candidates, we next sought to determine their neutralizing capacity and breadth of epitope recognition. Neutralizing antibodies were assessed against SARS-CoV-2 infection of Vero E6 cells *in vitro*. Although all adjuvanted groups elicited similarly high titers of RBD-binding IgG antibodies (10^5^-10^7^, **Suppl. Fig. S11a**), only RBD_surf_ neutralized the virus to a greater extent than unadjuvanted RBD_free_ at a plasma dilution of 10^−2.11^ (**Fig. 3a**). We then quantified the viral neutralization titer (VNT) as the dilution at which 50% of SARS-CoV-2-mediated cell death is neutralized. There was no significant difference between VNTs of human convalescent individual samples and RBD_surf_ plasma, and both groups induced higher VNTs compared to unadjuvanted RBD_free_ (**Fig. 3b**). Furthermore, the median VNT elicited by RBD_surf_ was 2.45, which falls within the FDA classification of “high titer” for convalescent plasma therapy (> 2.40)^41^. To ensure reproducibility, the neutralization assay was repeated with 3 different cohorts of n = 5 mice each, and no significant differences were observed between the resulting VNTs (**Suppl. Fig. S11b**).

**Figure 3.**
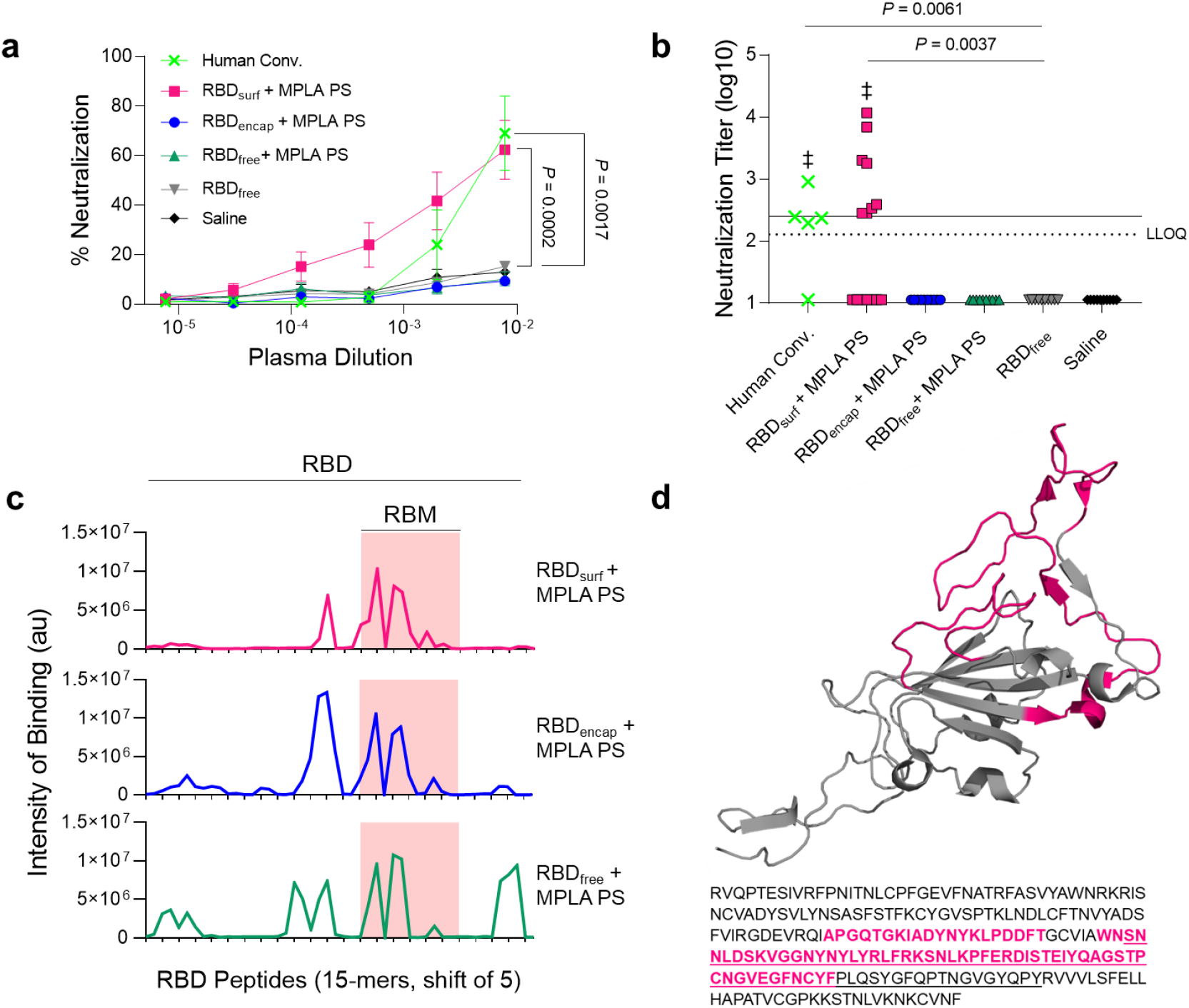
Antibodies induced by vaccination with RBD_surf_ + MPLA PS are neutralizing and localized to the receptor binding motif. **a**, Plasma from mice 1 week post-boost was tested for its ability to neutralize SARS-CoV-2 infection of Vero E6 cells *in vitro*. Percent neutralization for multiple plasma dilutions normalized to cells without virus (100%) or without plasma (0%). Data plotted as mean ± SEM for n = 5 convalescent human samples (human conv.) or 10-15 mice. Comparisons to unadjuvanted RBD_free_ were made for lowest dilution (10^−2.11^) using one-way ANOVA with Dunnett’s post-test. **b**, Viral neutralization titers, representing the plasma dilution at which 50% of SARS-CoV-2-mediated cell death is neutralized. Dashed line: lower limit of quantification (LLOQ = 2.11). For values below the LLOQ, LLOQ/2 values were plotted. Solid line: FDA recommendation for “high titer” classification (= 2.40). Comparisons were made using Kruskal-Wallis nonparametric test with Dunn’s post-test or Wilcoxon signed rank test (^‡^ ns, *P* > 0.05 compared to hypothetical value of 2.40). Symbols represent individual mice. **c**, Pooled plasma was then tested for antibody binding to linear epitopes using overlapping 15-amino-acid peptides with 5-amino-acid offsets, spanning the entire RBD sequence. X-axis represents sequential peptide number within the RBD amino acid sequence, and y-axis represents average luminescence for each peptide epitope. **d**, 3D structure of RBD with the receptor binding motif (RBM) underlined and main peptide sequences recognized by mouse plasma highlighted in pink (Protein Data Bank Entry 7DDD).

We next aimed to determine whether differences in neutralizing ability resulted from the epitope recognition breadth elicited by each vaccine formulation. To test this, we mapped the epitopes recognized by vaccine-elicited antibodies using a linear peptide array from the full-length RBD sequence. While IgG antibodies elicited by RBD_surf_ primarily recognized linear epitopes concentrated within the receptor binding motif of RBD (RBM; aa 438-508), RBD_encap_ and RBD_free_ + MPLA PS exhibited broader linear epitope diversity (**Fig. 3c,d, Suppl. Fig. S12**). Within RBD, the RBM is particularly critical for interacting with ACE-2, so antibodies specific for this region may have potent neutralizing potential^42,43^.

Because RBD_surf_ appeared to offer the advantage of improved neutralizing activity, while RBD_encap_ offered epitope diversity, we asked if co-administration would synergize to further enhance protection. To test this, we mixed RBD_surf_ and RBD_encap_ (with total RBD dose remaining constant) with MPLA PS and treated mice using the same vaccination schedule. As an additional control, we also investigated the humoral response of RBD_free_ adjuvanted with free MPLA. While both groups produced high RBD-specific IgG AUCs, neither exhibited neutralizing potential against the SARS-CoV-2 virus above background levels (**Suppl. Fig. S13a,b**). Analysis of the peptide arrays for these groups shows the presence of high-intensity-binding antibodies outside of the RBM (**Suppl. Fig. S13c**).

In summary, while all adjuvanted groups elicit high titers of RBD-specific antibodies, only RBD_surf_ generated neutralizing antibodies against SARS-CoV-2 at titers comparable to human convalescent plasma. Additionally, these antibodies uniquely bound to linear epitopes localized within the RBM, while all other groups produced antibodies with greater epitope breadth.

### All adjuvanted formulations increase T_fh_ and B cell activation in the dLN

Given the differences in antibody responses and neutralizing activity elicited by RBD_surf_ versus RBD_encap_ vaccination, we further investigated the phenotypes of the B and T cells involved in the initiation of the humoral immune response. All adjuvanted groups showed trends of higher frequencies of T follicular helper cells (T_fh_; CD4^+^CXCR5^+^BCL6^+^) in the injection site-draining lymph nodes (dLNs) 1 week post-boost compared to unadjuvanted RBD_free_ (**Fig. 4a, Suppl. Fig. S14**), although differences were only statistically significant for RBD_free_ + MPLA PS. Interestingly, a greater percentage of T_fh_ cells in all adjuvanted groups highly upregulated expression of ICOS, a co-stimulatory receptor important in T_fh_ activation and maintenance (**Fig. 4b**)^44^.

**Figure 4.**
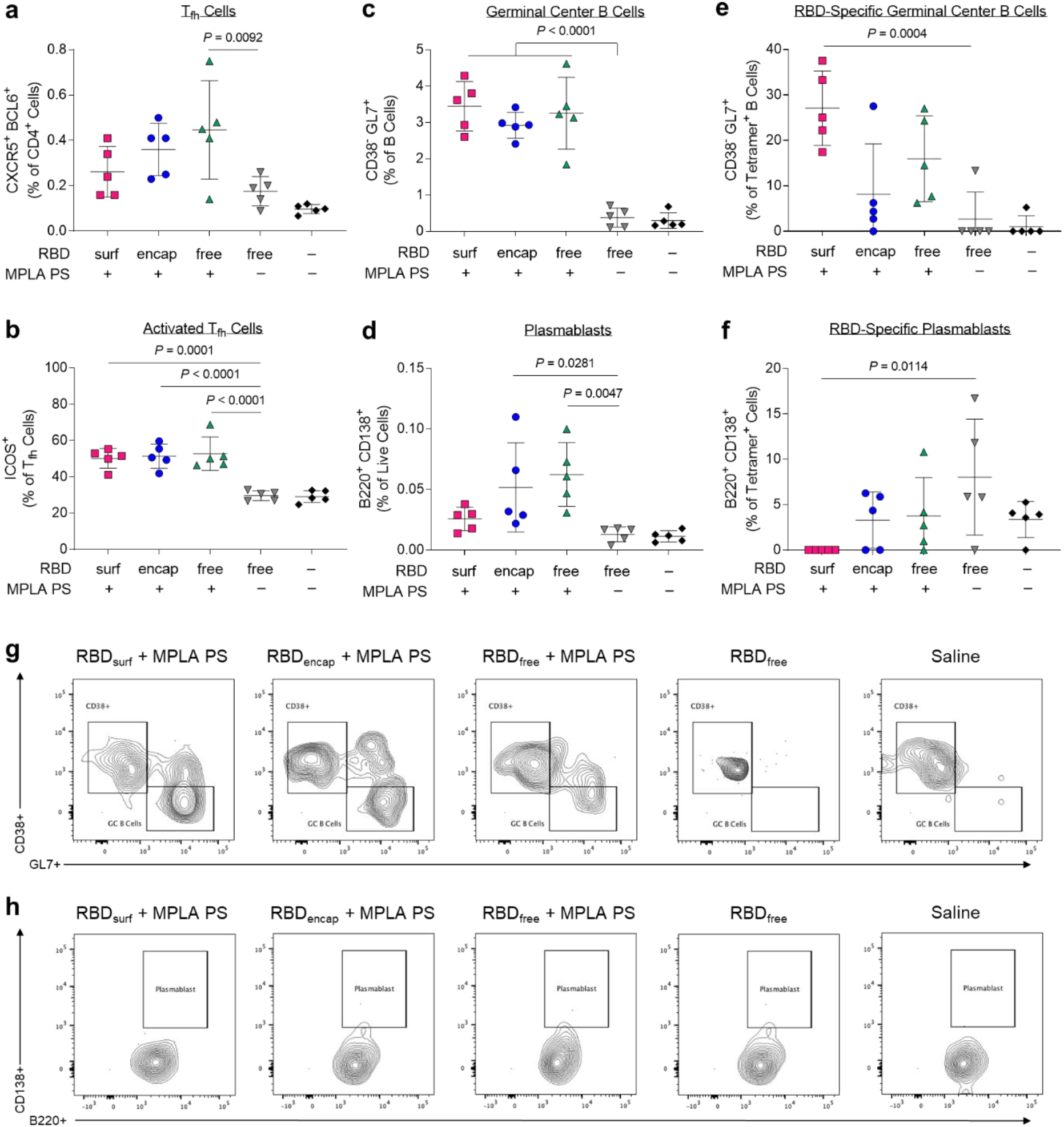
CD4^+^ T follicular helper cell (T_fh_) and B cells are activated by PS vaccine 1 week post-boost in the injection site-draining lymph nodes (dLN). **a**, T_fh_ cells (CXCR5^+^ BCL6^+^) of vaccinated mice quantified via flow cytometry as a percent of live CD4^+^ T cells. **b**, Highly activated ICOS^+^ T_fh_ cells quantified as percent of T_fh_ cells. **c**, Germinal center B cells (CD38^-^ GL7^+^) quantified as a percentage of total B cells (B220^+^ CD19^+^). **d**, Plasmablasts (B220^+^ CD138^+^) quantified as a percentage of total dLN cells. **e**, Germinal center B cells quantified as a percentage of RBD-specific B cells. **f**, Plasmablasts quantified as a percentage of RBD-specific dLN cells. **g-h**, Concatenated flow cytometry contour plots for n = 5 mice/group showing RBD-specific GC B cells (g) or plasmablasts (h). Data plotted as mean ± SD and represent 1 of 2 experiments with n = 5 mice each. Symbols represent individual mice. Comparisons to unadjuvanted RBD_free_ were made using one-way ANOVA with Dunn’s post-test.

Following activation by T_fh_ cells, naïve B cells can either undergo a germinal center (GC)-dependent response in which they become GC B cells and undergo cycles of class-switching and somatic hypermutation (SHM) before differentiation into long-lived plasma cells and class-switched memory B cells, or they can differentiate into short-lived plasmablasts or IgM memory cells in a GC-independent response^45^. A stronger GC response is valuable in vaccination because it results in higher affinity and longer-lived antibody production^46^. Overall frequencies of B cells (CD19^+^B220^+^) were unaffected across groups, but there were significantly lower frequencies of naïve IgD^+^ B cells in the adjuvanted groups compared to unadjuvanted RBD_free_ (**Suppl. Figs. S15, S16**). All adjuvanted groups generated GC responses, characterized by increased frequencies of GC B cells (CD38^-^GL7^+^) in the dLN compared to unadjuvanted RBD_free_ (**Fig. 4c**). Both RBD_encap_ and RBD_free_ + MPLA PS formulations, but not RBD_surf,_ significantly increased the frequencies of plasmablasts (B220^+^CD138^+^) in the dLN compared to unadjuvanted RBD_free_ (**Fig. 4d**).

To determine the antigen-specificity of the B cells, we developed a set of fluorescent RBD protein tetrameric probes. To ensure selectivity, B cells were considered RBD-specific if they were double-positive for both PE- and APC-conjugated RBD-tetramers and negative for the non-specific control protein tetramer (**Suppl. Fig. S17**). RBD_free_ + MPLA PS was the only formulation to significantly increase the frequency of RBD-specific B cells in the dLN (**Suppl. Fig. S18**). We next sought to further determine the phenotype of these RBD-specific B cells. RBD_surf,_ unlike the other adjuvanted formulations, led to a significantly higher fraction of RBD-specific B cells with GC B cell phenotype, suggesting a more robust GC response to RBD (**Fig. 4e**). RBD_surf_ was also the only formulation with a significantly lower fraction of plasmablasts within the RBD-specific B cell subset compared to unadjuvanted RBD_free_ (**Fig. 4f**). These differences are also visually apparent in pooled flow cytometry plots for RBD-specific GC B cells (**Fig. 4g**) and plasmablasts (**Fig. 4h**). In summary, all adjuvanted formulations of RBD increased activation of T_fh_ cells and GC B cells in the dLN, but within the RBD-specific B cell population, only RBD_surf_ generated a higher fraction of GC B cells and a lower fraction of plasmablasts.

### Vaccination with polymersome-formulated RBD generates RBD-specific Th1 T cell responses

Having demonstrated that our PS vaccines can generate strong humoral responses, we next sought to determine their capacity to generate robust CD8^+^ and CD4^+^ T cell immunity. In order to assess the RBD-specific T cell response, we isolated cells from the dLNs of vaccinated mice 1 week post-boost. Prior to intracellular staining, cells were restimulated with RBD peptide pools for 6 hours. The RBD-specific response was quantified by subtracting the signal from cells incubated in media alone from those incubated with RBD peptide pools (**Suppl. Fig. S19**). Only PS-formulated RBD groups RBD_surf_ and RBD_encap_ generated significantly higher frequencies of IFNγ^+^, bifunctional IFNγ^+^TNFα^+^, and IL-2^+^ secreting CD8^+^ T cells compared to unadjuvanted RBD_free_ (**Fig. 5a**). Similar trends were seen in the CD4^+^ T cell compartment. Treatment with RBD_surf_ and RBD_encap_ but not RBD_free_ + MPLA PS led to significantly higher frequencies of IFNγ^+^ and IL-2^+^ secreting CD4^+^ T cells, although the increase in bifunctional IFN γ^+^TNFα^+^ was not statistically significant (**Fig. 5b**). The production of these cytokines implies a Th1 T cell response, which is correlated with less severe cases of SARS-CoV infection^21^.

**Figure 5.**
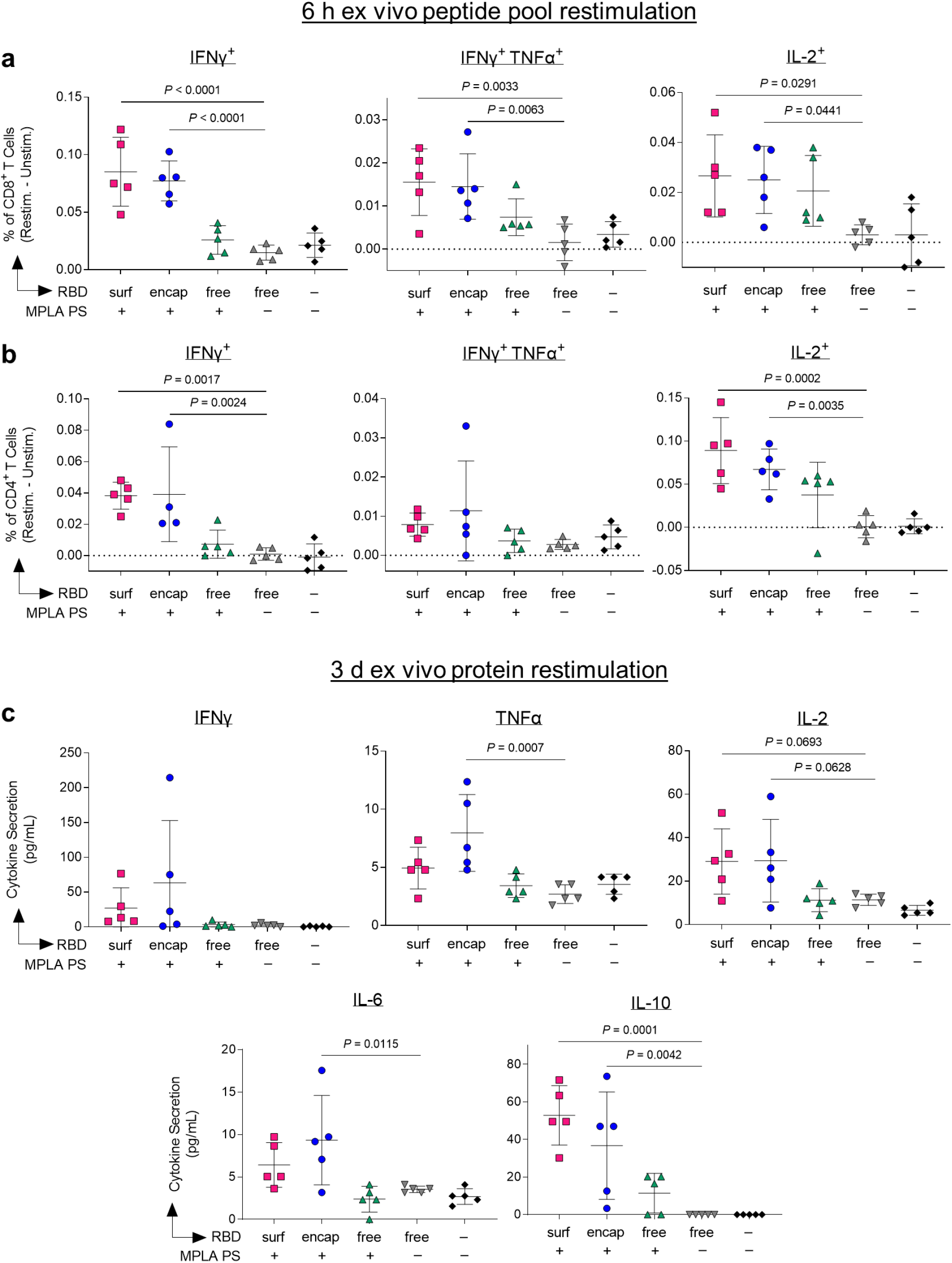
Vaccination with polymersome-formulated RBD improves antigen-specific T cell responses in mice. Cells isolated from the injection-site draining lymph nodes of PS-vaccinated mice 1 week post-boost were restimulated ex vivo for 6 h with RBD peptide pools or full RBD protein and analyzed by flow cytometry or multiplexed ELISA, respectively. **a-b**, percentages of cytokine-positive CD8^+^ T cells (a) and CD4^+^ T cells (b) in response to RBD peptide pools, subtracting unstimulated controls. **c**, Pro-inflammatory cytokine levels released into the supernatant measured after 3 d restimulation with whole RBD protein. Data plotted as mean ± SD and represent 1 of 2 experiments with n = 5 mice each. Symbols represent individual mice. Comparisons to unadjuvanted RBD_free_ were made using one-way ANOVA with Dunn’s post-test.

For further validation of the cytokine response, cells from the dLN isolated from the same vaccinated mice were restimulated with full RBD protein *ex vivo* for 3 days, followed by quantification of cytokines in the supernatant. The RBD-specific response was quantified by subtracting unstimulated signal from stimulated signal as above. The levels of Th1-type cytokines detected were consistent with intracellular staining data. Levels of IFNγ and IL-2 were modestly but not significantly increased in the RBD_surf_ and RBD_encap_ groups compared to the RBD_free_ group, while cells from RBD_encap_-treated mice produced TNFα at significantly higher levels than unadjuvanted RBD_free_ (**Fig. 5c**). RBD_encap_ also led to increased production of IL-6 and IL-10, which are pleiotropic cytokines known to be secreted during Th1 responses (**Fig. 5c**)^47,48^. Levels of secreted Th2-type cytokines were also measured. More IL-4 was produced in the supernatant of samples treated with RBD_surf_ and RBD_encap_ compared to RBD_free_, albeit at an overall low concentration. There was no significant elevation of IL-5 secretion across any of the treatment groups compared to the saline control and no detectable levels of IL-13 in any sample (**Suppl. Fig. S20)**. In summary, vaccination with RBD delivered via polymersome formulations generated stronger RBD-specific Th1-type CD8^+^ and CD4^+^ T cell responses than unadjuvanted RBD_free_, while RBD_free_ + MPLA PS did not.

## DISCUSSION

In this study, we developed antigen-decorated, oxidation-sensitive polymersomes that mimic virus particles as next-generation nanovaccines. While all adjuvanted formulations generated long-lived RBD-binding IgG responses, surface conjugation of antigen was necessary to generate neutralizing antibodies against SARS-CoV-2. More generally, this comparison of antigen formulation on the quality of immune response offers valuable insight into vaccine design, demonstrating the benefits of surface antigen display.

The differences in the immune responses elicited by the two PS antigen formulations, RBD_surf_ and RBD_encap_, suggest that surface display of antigen leads to stronger GC responses, while PS-encapsulated antigen elicits more predominantly an extrafollicular response. Though RBD-specific GC B cells are present in the dLN after treatment with both formulations, a much higher percentage of the RBD-specific B cells recovered after vaccination with RBD_surf_ exhibited a GC B cell phenotype. This higher percentage could be due to the induction of higher numbers of RBD-specific GC B cells after vaccination with RBD_surf_ or an increase in their affinity, leading to easier detection via RBD protein tetramer staining. Both possibilities suggest a more robust GC response, as GC responses are necessary for an efficient SHM leading to increased B cell affinity^49^. Evidence for affinity maturation due to SHM also includes the relatively few epitopes on the RBD linear peptide array to which IgG from RBD_surf_-treated groups were specific compared to the other adjuvanted groups. Clonal bursts in GC responses can lead to rapid expansion of high-affinity SHM variants and loss of overall clonal diversity^50^. A difference in the affinity of RBD-specific IgG generated by these vaccines may also explain the neutralization ability of the plasma after vaccination with RBD_surf_ but not RBD_encap_. Further data that suggest that RBD_encap_ elicits a more extrafollicular response include the fast initial antibody response after priming by RBD_encap_, which resulted in RBD-specific IgG AUC 1 week after the initial prime that was significantly higher than that of RBD_surf_. In an extrafollicular B cell response, B cells can differentiate immediately into plasmablasts and begin secreting antibodies after initial T cell help, whereas B cells that enter GCs delay the antibody response by several days^45^. The preferential differentiation into plasmablasts rather than GC B cells was also evident in the increased fraction of plasmablasts within the RBD-specific B cell population generated by RBD_encap_. In vaccine development, the generation of robust GC reactions is preferable to extrafollicular responses because GC formation will usually result in higher affinity B cells, increased memory B cells and increased long-lived plasma cells in the bone marrow.^45^

The difference in mechanism of B cell activation by these two RBD formulations may be explained by previous studies on multimerization of antigen and use of virus-like particles in vaccination. Virus-like particles are multi-protein supra-molecular structures constructed of many identical protein copies. Their multimeric nature has been associated with the induction of potent antibody responses due to BCR crosslinking in the presence of CD4^+^ T cell help^51^. Mechanistic studies by Kato et al. demonstrated that increasing antigen valency can enhance the early activation and proliferation of antigen-specific B cells as well as increase B cell accumulation at the T-B border leading to increased differentiation of antigen-specific B cells into GC B cell and plasma cells^52^. Furthermore, in contrast to RBD_encap_, RBD_surf_ most likely efficiently exposes for BCR interaction the conformational epitopes of RBD reported to be targeted by neutralizing antibodies in plasma of convalescent or vaccinated individuals^7–10^. Thus, the multimerization of RBD on the PS surface in addition to its increased availability to B cells led to an improved functional quality of humoral response compared to encapsulated or unformulated RBD.

The type of immune response to SARS-CoV-2 may have important implications in how the infection is cleared and should be considered in vaccine design^21^. Less severe cases of the original SARS-CoV were associated with increased Th1-type cell responses^53,54^. In contrast, Th2-type responses were associated with increased pathology due to antibody-dependent enhancement, and several vaccine formulations against SARS-CoV tested in animal models showed signs of immunopathology due to Th2 cell-mediated eosinophil infiltration^55,56^. In this study, we measured the ratio of IgG1/IgG2b and the cytokine profile of dLN cells restimulated with RBD protein in order to characterize the type of immune responses generated by our vaccine formulations. RBD_encap_ and RBD_free_ + MPLA PS generated IgG2b/IgG1 ratios around 1, suggesting a balance between type 1 and type 2 immunity, whereas RBD_surf_ generated a response with a slightly IgG1-bias, suggestive of a Th2-skewed response. However, both RBD_surf_ and RBD_encapf_ generated significant levels of the Th1-type cytokines IFNγ, TNFα and IL-2. This Th1-type response is likely induced by the MPLA PS, as MPLA has been shown to generate strong type-1 immunity^57^. Although RBD_surf_ generated a slightly IgG1-biased antibody response, it did not result in an increase in Th2-type cytokines upon restimulation compared to RBD_encap._ Combined with the evidence of Th1-type cytokine production upon restimulation, we believe the risk of adverse events related to Th2-type responses is low.

The PS-formulated RBD vaccines were able to generate stronger Th1-driven CD4^+^ and CD8^+^ T cell responses compared to RBD_free_. We previously demonstrated that as an antigen delivery vehicle, PEG-PPS polymersomes could improve cross-presentation to CD8^+^ T cells^33^. This increased T cell response likely occurs because of both enhanced APC targeting as well as rapid endosomal antigen release^33^. APCs are able to accumulate membrane-impermeable nanocarriers such as PS more efficiently than other cell types due to their constitutive macropinocytosis^58^. Once endocytosed, PEG-PPS polymersomes require only a small amount of oxidation to release encapsulated antigen, and payload delivery to the cytosol is not restricted to endosomal compartments with reductive or acidic conditions^33^. Unlike antigen-encapsulated PS, however, acidification may be important for proteolytic degradation of RBD_surf_ and translocation to the cytosol, as evidenced by studies on VLPs^59,60^. Additionally, peptides derived from large antigen particles have been found to enter the cross‐presentation pathways more efficiently than those derived from soluble antigens, which may provide rationale for the enhanced CD8^+^ response of RBD_surf_, whereas no such benefit was seen for adjuvanted RBD_free_^61^. Therefore, antigen formulation using PS to improve T cell responses could be beneficial in the engineering of future vaccines against cancer or other infectious diseases for which T cell immune responses are thought to be necessary for protection, such as herpesviruses, human immunodeficiency virus, and hepatitis C virus^62^.

In summary, we have demonstrated that a polymersome-based antigen and adjuvant delivery system generates robust humoral immunity and neutralizing antibody titers, as well as T cell responses, against a key SARS-CoV-2 vaccine target, the RBD of Spike. This platform technology is amenable to a wide variety of antigens and formulated or soluble adjuvants. Once the type and dose of adjuvant has been optimized for a given application, a single particle could be used to deliver both antigen and adjuvant to APCs in the injection site-draining lymph node. Additionally, multiple antigens, for example from different viruses or different strains or variants of the same virus, could be conjugated to the same particle as a strategy to induce cross-reactive neutralizing antibodies^20^. Importantly, both surface-decorated and antigen-encapsulated polymersomes remained stable at 4 °C for at least 6 months, as indicated by consistent particle size and absence of antigen released into solution. Vaccines that exhibit long-term stability without requiring sub-zero temperatures will likely be important for widespread vaccine distribution, for example to rural populations or developing nations with poor cold chain network. The evaluation of RBD-decorated polymersomes presented here could thus provide insight into the next generation of stable formulations of nanovaccines to combat the current COVID-19 pandemic as well as future viral outbreaks.

## METHODS

### RBD production and purification

For production of the Spike protein RBD (Spike_319-541_; GenBank: MN908947.3), we obtained expression plasmids on pCAGGS backbone containing mammalian codon-optimized sequences for this gene from Florian Krammer’s laboratory (Icahn School of Medicine at Mount Sinai, New York, NY)^63^. Suspension-adapted HEK-293F cells were maintained in serum-free Free Style 293 Expression Medium (Gibco). On the day of transfection, cells were inoculated into at a concentration of 1 × 10^6^ cells mL^-1^. Plasmid DNA (1 mg mL^-1^) was mixed with linear 25 kDa polyethyleneimine (2 mg mL^-1^; Polysciences) and co-transfected in OptiPRO SFM medium (4% final concentration; Thermo Fisher). Flasks were cultured in an orbital shaking incubator (135 rpm, 37 °C, 5% CO_2_) for 7 days. Culture medium was then collected by centrifugation, filtered, and loaded into a HisTrap HP 5 mL column (GE Healthcare) using an ÄKTA pure 25 (GE Healthcare). After washing the column with wash buffer (20 mM NaH_2_PO_4_ and 0.5 M NaCl, pH 8.0), protein was eluted using a gradient of 500 mM imidazole in wash buffer. The protein was further purified by size-exclusion chromatography using a HiLoad Superdex 200PG column (GE Healthcare) with PBS as an eluent. Dimers of RBD were reduced by the addition of dithiothreitol (1 mM) which was subsequently dialyzed against PBS. All purification steps were carried out at 4 °C. The expressed proteins were verified to be >90% pure through SDS–PAGE. The purified proteins were tested for endotoxin using a HEK-Blue™ TLR4 reporter cell line (InvivoGen), and the endotoxin levels were confirmed to be below 0.01 EU mL^−1^. Protein concentration was determined by absorbance at 280 nm using a NanoDrop spectrophotometer (Thermo Scientific). Proteins were stored at a concentration of 4 mg mL^-1^ at −80 °C until use.

### PS formulation

PEG-PPS polymersomes were formulated by thin film rehydration as previously described^33^. In brief, 20 mg of polymer was dissolved in 750 µL dichloromethane (DCM), and DCM was removed by vacuum desiccation overnight. 1 mL of PBS was then added to the vial, which was rotated at room temperature (RT) for 24 h to allow complete dispersal of the polymer. The solution was then sequentially extruded through 0.8, 0.4, 0.2, and 0.1 µm pore membranes (Whatman). To formulate RBD-encapsulated PS, 250 µL of PBS containing 4 mg/mL RBD was added to the polymer thin film for rehydration, and the solution was rotated at 4 °C for 72 h before extrusion as above. After extrusion, RBD-encapsulated polymersomes were passed through a sepharaose size exclusion chromatography (SEC) column to remove unencapsulated free RBD. The RBD content was quantified by SDS-PAGE using mini-protein TGX stain-free precast gels (Bio-Rad). Gels were imaged on a ChemiDoc XRS+ Gel Documentation System (Bio-Rad) and analyzed using ImageJ.

RBD-surface-conjugated PS were synthesized by first formulating empty PS as above consisting of 25% N_3_-PEG-PPS. RBD was conjugated to a *sulfo* DBCO-Maleimide linker (Click Chemistry Tools) at the molar ratio of 1:5 (RBD:linker) and reacted for 2.5 hours. Unconjugated linker was removed by Zeba spin desalting columns (7K MWCO; ThermoFisher). The resulting RBD-linker was analyzed by MALDI-TOF MS using an α-cyano-4-hydroxycinnamic acid (Sigma-Aldrich) matrix and a Bruker ultrafleXtreme MALDI TOF/TOF instrument. RBD-linker (4 wt% of polymersome) was then incubated with empty N_3_-PEG-PPS polymersomes overnight at RT. RBD-surface-conjugated PS were then passed through SEC to remove unconjugated RBD-linker. The conjugation was monitored by SDS-PAGE, and the final RBD content was quantified by the CBQCA Protein Quantitation Kit (ThermoFisher). Fluorescently labeled PS were prepared by conjugating AlexaFluor-647 alkyne dye (ThermoFisher) to PS containing 25% N_3_-PEG-PPS. The dye was mixed with PS at a molar ratio of 1:20 (dye:N_3_-PEG-PPS), and the solution was stirred overnight at RT. Labeled PS were then passed through SEC to remove unconjugated dye. To make the fluorescently labeled RBD_surf_, labeled PS were prepared as above and conjugated with RBD as described, followed by SEC purification. Final RBD content was quantified by the CBQCA Protein Quantitation Kit (ThermoFisher).

MPLA PS were fabricated by flash nanoprecipitation using a 3D printed impingement jets mixer^64^. 20 mg PEG-PPS and 2 mg MPLA (PHAD®; Avanti Polar Lipids) were dissolved in 500 µL tetrahydrofuran (THF) and loaded into a 1 mL plastic disposable syringe. 500 µL PBS was loaded into a second syringe, and the two solutions were impinged against one another slowly within the mixer by hand. The impinged solution was immediately vortexed to form a homogenous polymersome solution which was then extruded and purified by SEC as described above. MPLA loading was quantified using a liquid chromatography-tandem mass spectrometry (LC-MS/MS) method as previously described^65^ using PHAD®-504 (Avanti Polar Lipids) as an internal standard on an Agilent 6460 Triple Quad MS-MS with 1290 UHPLC.

### PS characterization

The size and polydispersity index (PDI) of all the polymersome formulations were measured by dynamic light scattering (DLS) using a Zetasizer Nano ZS90 (Malvern Instruments). Cryogenic electron microscopy (cryoEM) images were obtained on a FEI Talos 200kV cryoEM dedicated electron microscope.

### MPLA PS *in vitro* activity

To determine TLR4 activation, HEK-Blue™ TLR4 cells (Invivogen) were incubated with increasing concentrations of MPLA PS for 24 h 37 °C in a 5% CO_2_ incubator. NF-kB-induced SEAP activity was detected using QUANTI-Blue™ (Invivogen) and by reading the OD at 650 nm. For dendritic cell activation experiments, BMDCs were prepared from C57Bl/6 mice (Jackson) as previously described.^66^ After 9 days of culture, cells were seeded at 2 × 10^5^ cells/well in round-bottom 96-well plates (Fisher Scientific) in IMDM with 10% FBS and 2% penicillin/streptomycin (Life Technologies). Cells were treated with varying concentrations of MPLA PS or free MPLA and incubated for 24 h at 37 °C in a 5% CO_2_ incubator. After 24 h, the supernatant was collected, and cytokine concentration was measured using a multiplexed mouse Th cytokine panel (BioLegend) according to the manufacturer’s instructions.

### RBD_surf_ *in vitro* activity

For the cell-based hACE2-binding assay, human embryonic kidney (HEK)-293T cells overexpressing human ACE-2 (HEK-hACE2) were obtained from BEI Resources (NIH NIAID). Fluorescently labeled empty PS, RBD_surf_, or RBD_free_ was incubated at varying concentrations with 5 × 10^4^ HEK-hACE2 cells at 4 °C for 20 min. Cells were then washed three times in PBS with 2% FBS. Binding of RBD to hACE-2 on the cell surface was assessed via the mean fluorescent intensity measured by flow cytometry using BD LSRFortessa (BD Biosciences).

### Production of RBD protein tetramers

RBD protein expressed with AviTag was purchased from GenScript. Site-specific biotinylation of the AviTag was performed using BirA Biotin-Protein Ligase Reaction kit (Avidity). Next, unconjugated biotin was removed using Zeba spin desalting columns, 7K MWCO (ThermoFisher). The quantification of reacted biotin was performed using the Pierce Biotin Quantification Kit (ThermoFisher). Biotinylated RBD was incubated with either streptavidin-conjugated PE (Biolegend) or streptavidin-conjugated APC fluorophores (Biolegend) for 20 min on ice at a molar ratio of 4:1 of biotin to streptavidin. Streptavidin-conjugated FITC (BioLegend) was reacted with excess free biotin to form a non-RBD-specific streptavidin probe as a control. Tetramer formation was confirmed using SDS-PAGE gel. Cells were stained for flow cytometry with all three streptavidin probes at the same time as other fluorescent surface markers at a volumetric ratio of 1:100 for RBD-streptavidin-PE and 1:200 for RBD-streptavidin-APC and biotin-streptavidin-FITC.

### Mouse vaccination experiments

All experiments were performed in accordance with the Institutional Animal Care and Use Committee at the University of Chicago. Female 8-week-old C57BL/6 mice (Jackson Laboratory) were randomly assigned to cohorts of n = 5 and vaccinated with 10 μg of antigen and 5 μg of adjuvant s.c. in the hock (either 2 or 4 hocks) and boosted on day 21. On day 7, 14, 20, and 28 post vaccination, 100 μL of blood was collected in EDTA-K2-coated tubes (Milian), and plasma was separated by centrifugation and stored at −80 °C until use. On day 28 after initial vaccination, mice were sacrificed. Splenocytes and lymph node cells from draining lymph nodes were collected. Single-cell suspensions of the lymph node were prepared by digestion in collagenase D for 45 min at 37 °C. Splenocytes and lymph node cells were filtered through a 70 μm cell strainer. Splenocytes were then incubated in ACK lysis buffer to remove red blood cells. Lymph node cells were stained for T_fh_ cells and RBD-specific B cells using fluorescent probes listed in Supplementary Tables 1 and 2, respectively. Samples were acquired on a BD LSR-Fortessa (BD) and analyzed using FlowJo™ software. Representative gating strategies used to identify the cell populations are shown in Supplemental Figures 14 (T_fh_ cells) and 15 (RBD-specific B cells).

To assess antigen-specific cytokine production by T cells, 1 × 10^6^ lymph node cells were incubated with pools of 15-mer peptides overlapping by 10 amino acids covering the N-terminus of SARS-CoV-2 Spike protein up to the furin cleavage site (S1 pool; PepMix SARS-CoV-2 Spike Glycoprotein, JPT) for 6 h at 37 °C with 5% CO_2_. Monensin (GolgiStop, BD) was added after 2 h of incubation to inhibit cytokine secretion. Cells were stained for surface markers using fluorescent monoclonal antibodies (mAbs). Cells were subsequently fixed and permeabilized using BD Cytofix/Cytoperm™, and intracellular cytokines were stained using fluorescent-mAbs listed in Supplementary Table 3. Samples were acquired on a BD LSR-Fortessa (BD) and analyzed using FlowJo™ software. A representative gating strategy used to identify the cell populations is shown in Supplementary Figure 19. To assess antigen-specific cytokine secretion, lymph node cells were plated at 5 × 10^5^ cells/well were incubated with 100 μg RBD for 3 d at 37 °C with 5% CO_2_. After 3 d, the supernatant was analyzed for cytokine concentration via a multiplexed mouse Th cytokine panel (BioLegend) according to the manufacturer’s instructions. Samples were aquired on an Attune NxT flow cytometer (ThermoFisher), and analyzed with LEGENDplex v8.0 software.

For long-term experiments, mice were not sacrificed but continued to be bled on day 2, 7, 14 and then every 2 weeks for 80 days.

### RBD-binding ELISA

Plasma was assessed for anti-RBD IgG and IgA by ELISA. 96-well ELISA plates (Costar high-bind flat-bottom plates, Corning) were coated with 10 μg/mL RBD in carbonate buffer (50 mM sodium carbonate/sodium bicarbonate, pH 9.6) overnight at 4 °C. The following day, plates were washed three times in PBS with 0.05% Tween 20 (PBS-T) and then blocked with 1x casein (Sigma) for 1 h at RT. Following blocking, wells were washed three times with PBS-T and further incubated with six 10-fold dilutions of plasma in 1x casein for 2 h at RT. Wells were then washed five times with PBS-T and incubated for an additional hour at RT with horseradish peroxide (HRP)-conjugated antibodies against mouse IgG, IgG1, IgG2b, IgG3, or IgA (Southern Biotech). After five washes with PBS-T, bound RBD-specific Ig was detected with tetramethylbenzidine (TMB) substrate. Stop solution (3% H_2_SO_4_ + 1% HCl) was added after 18 min of TMB incubation at RT, and the OD was measured at 450 and 570 nm on an Epoch Microplate Spectrophotometer (BioTek). Background signal at 570 nm was subtracted from the OD at 450 nm. Fold-change over the average of blank wells was then calculated and log-transformed. The area under the curve (AUC) of log-transformed fold change versus log-transformed dilution was then calculated.

### RBD-binding IgG ELISpot assay

ELISpot plates (Millipore IP Filter plate) were coated with 20 µg/mL RBD in sterile PBS overnight at 4 °C. Plates were then blocked using ELISpot Media (RPMI 1640, 1% glutamine, 10% fetal bovine serum, 1% penicillin-streptomycin) for 2 hours at 37 °C. Splenocytes from vaccinated mice were seeded in triplicate at a starting concentration of 6.75 × 10^5^ cell/well and diluted seven times in 3-fold serial dilutions. Plates were incubated for 18 hours at 37 °C and 5% CO_2_ after which the cells were washed five times in PBS. Wells were incubated with 100 µL IgG-biotin HU adsorbed (Southern Biotech) for 2 h at RT. Next, plates were washed four times in PBS followed by the addition of 100 µL HRP-conjugated streptavidin/well for 1 h at RT. Plates were washed again and incubated with 100 µL TMB/well for 5 minutes until distinct spots emerged. Finally, plates are then washed three times with distilled water and left to dry completely in a laminar flow hood. A CTL ImmunoSpot Analyzer was used to image plates, count spots, and perform quality control.

### SARS-CoV-2 virus neutralization assay

SARS-CoV-2 viruses (400 plaque forming units; strain nCov/Washington/1/2020, provided by the National Biocontainment Laboratory, Galveston TX, USA) were incubated with 4-fold serial dilutions of heat-inactivated plasma from vaccinated or control mice and for 1 h at 37 °C in DMEM with 2% fetal bovine serum, penicillin-streptomycin (100 U/mL penicillin, 100 μg/mL streptomycin), and non-essential amino acids (10 mM, glycine, L-alanine, L-asparagine, L-aspartic acid, L-glutamic acid, L-proline, L-serine; Gibco). The pre-incubated viruses were then applied to Vero-E6 cell monolayers, and the cells were maintained until > 90% cell death for the negative control (4-5 d). Cells were then washed with PBS, fixed with 10% formalin, stained with crystal violet, and quantified with a Tecan infinite m200 microplate reader (excitation/emission 592 nm/636 nm). Neutralization titer is measured as the greatest dilution that inhibits 50% of SARS-CoV-2 induced cell death (EC50). To determine the EC50, data were fit using a least squares variable slope four-parameter model. To ensure realistic EC50 values, we considered a dilution (1/X) of X = 10^−1^ to be 100% neutralizing and a dilution of X = 10^8^ to be 0% neutralizing and constrained EC50 > 0. Plasma from convalescent human COVID-19 patients were provided by Ali Ellebedy (Washington University School of Medicine, St. Louis, MO; Catalog # NR-53661, NR-53662, NR-53663, NR-53664, and NR-53665).

### Peptide array analysis

Antibody specificity to linear epitopes of the spike protein was analyzed using a CelluSpots™ Covid19_hullB Peptide Array (Intavis Peptide Services, Tubingen, Germany) according to the manufacturer’s protocol. The array comprises 254 peptides spanning the full-length sequence of the Spike protein (NCBI GenBank accession # QHD43416.1), with each 15-mer peptide offset from the previous one by 5 amino acids. Briefly, peptide arrays were blocked in 1x casein solution at 4 °C overnight. Arrays were then incubated with pooled serum diluted 1:200 in 1x casein for 6 h at RT on an orbital shaker (60 rpm). After 6 h, arrays were washed four times with PBS-T and incubated for an addition 2 h at RT, 60 rpm with goat anti-mouse IgG conjugated to HRP (Southern Biotech) diluted 1:5000 in 1x casein. Arrays were washed another four times with PBS-T. Spots were detected with Clarity™ Western ECL Substrate (Bio-Rad), and chemiluminescence was measured using a ChemiDoc XRS+ Gel Documentation System (Bio-Rad). Spots were analyzed using Spotfinder software (version v3.2.1).

### Software packages and statistical analysis

Statistical analysis was performed using GraphPad Prism 9 (GraphPad). Data were analyzed using one-way ANOVA with Dunn’s or Tukey’s post-hoc correction for multiple hypothesis testing unless otherwise stated. All flow cytometry data were analyzed using FlowJo_v10.7.2 software (FlowJo LLC, BD Biosciences). Figures 1 and 2 were created using BioRender (https://biorender.com) as part of an Academic License through the Chicago Immunoengineering Innovation Center.

## ACKNOWLEDGMENTS

We are grateful for funding support through a pilot project grant from the Chicago Immunoengineering Innovation Center of the University of Chicago, as well as the Chicago Biomedical Consortium COVID-19 Response Award (# CR-002) to M.A.S. In addition, a number of researchers were supported by fellowships via the NIH NHLBI (# T32-HL007605 to L.R.V.), Canadian Institutes of Health Research (#201910MFE-430736-73744 to N.M.), University of Chicago Comprehensive Cancer Center (Sigal Fellowship in Immuno-Oncology to N.M.), NIH NCI (# F30-CA221250 to M.R.S.), NIH NIGMS (# T32-GM007281 to M.R.S.) and NIH NIAID (# T32-AI007090 to T.M.M. and A.C.T.). We are grateful to the laboratory of Florian Krammer (Icahn School of Medicine at Mount Sinai, New York City, NY) for providing plasmids coding for the Spike RBD, produced with support from the NIH NIAID (Contract # HHSN272201400008C). We are also grateful to the groups of Jesse Bloom (Fred Hutchinson Cancer Research Center, Seattle, WA) and Ali Ellebedy (Washington University School of Medicine, St. Louis, MO) for contributing reagents via the NIH NIAID BEI Resources repository. We acknowledge helpful discussions with Patrick C. Wilson, Jenna J. Guthmiller, Anne I. Sperling, and Aaron Esser-Kahn (University of Chicago, Chicago, IL) and with Robert Baker and David Boltz (Illinois Institute of Technology Research Institute, Chicago, IL) that were instrumental to experimental planning and model development. We acknowledge Suzana Gomes and Tera Lavoie for technical assistance. Parts of this work were carried out at the Cytometry and Antibody Technology Core Facility (Cancer Center Support Grant P30CA014599), the Soft Matter Characterization Facility, the Mass Spectrometry Facility (NSF instrumentation grant CHE-1048528), the Nuclear Magnetic Resonance Facility, the Advanced Electron Microscopy Facility (RRID:SCR_019198), and the Human Immunologic Monitoring Facility (RRID:SCR_017916) at the University of Chicago.

## SUPPLEMENTARY INFORMATION

**Supplementary Figure S1.**
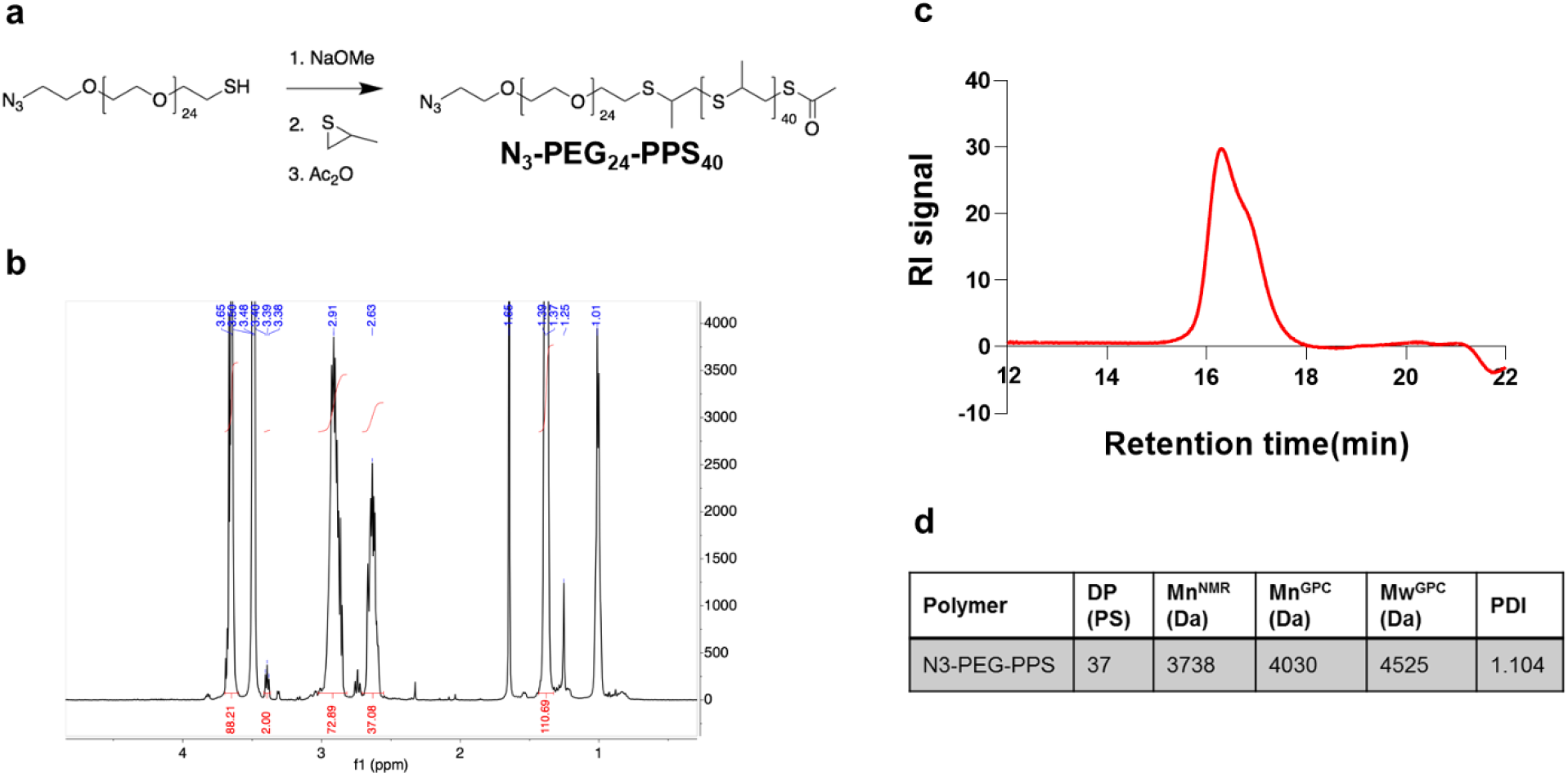
Synthesis and characterization of N_3_-PEG-PPS. **a**, Synthetic route, **b**, ^1^H NMR spectrum, **c**, gel permeation chromatography (GPC) trace, and **d**, summary of physiochemical properties of N_3_-PEG-PPS.

**Supplementary Figure S2.**
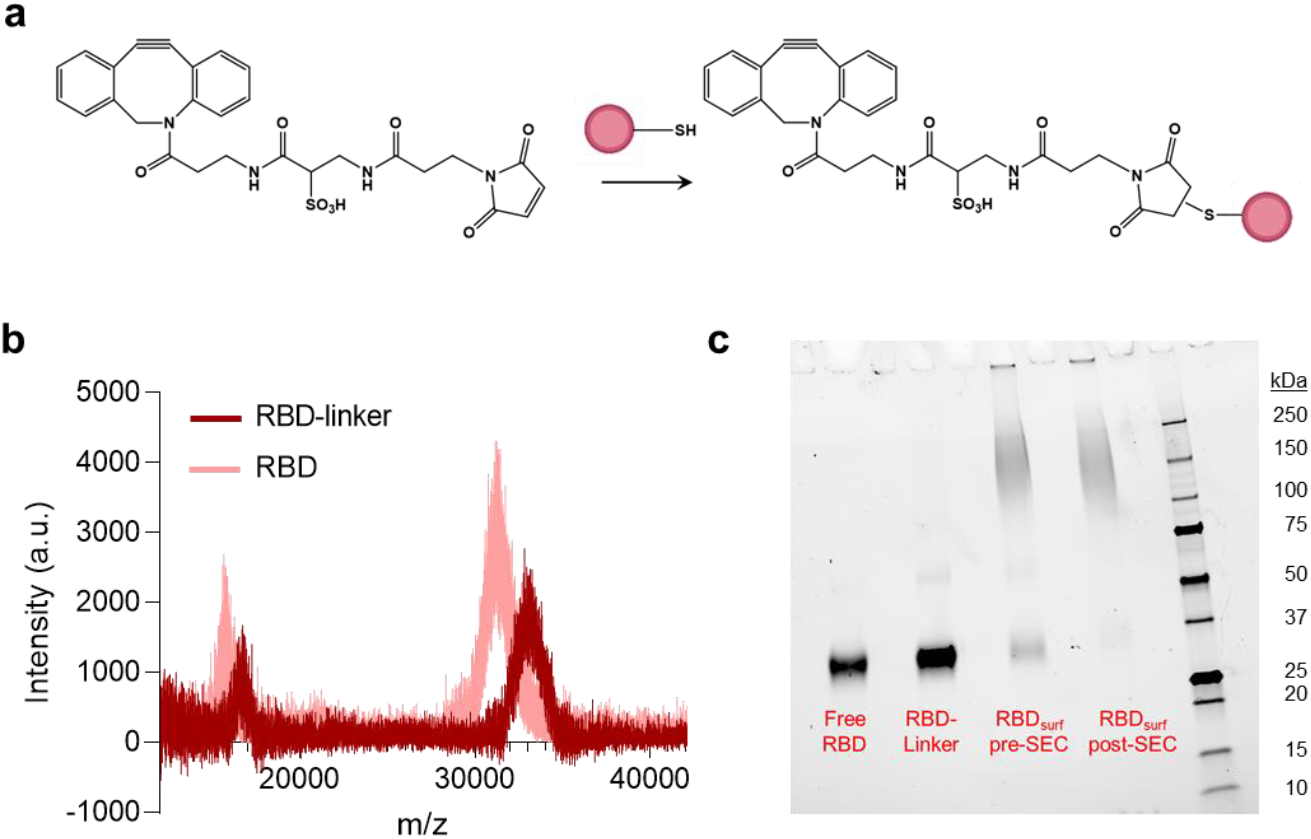
Synthesis and characterization of RBD-linker. **a**, Synthetic route, **b**, MALDI of RBD-linker and free RBD, **c**, SDS PAGE of free RBD, RBD-linker, RBD_surf_ before size exclusion chromatography (SEC) and purified RBD_surf_ post-SEC.

**Supplementary Figure S3.**
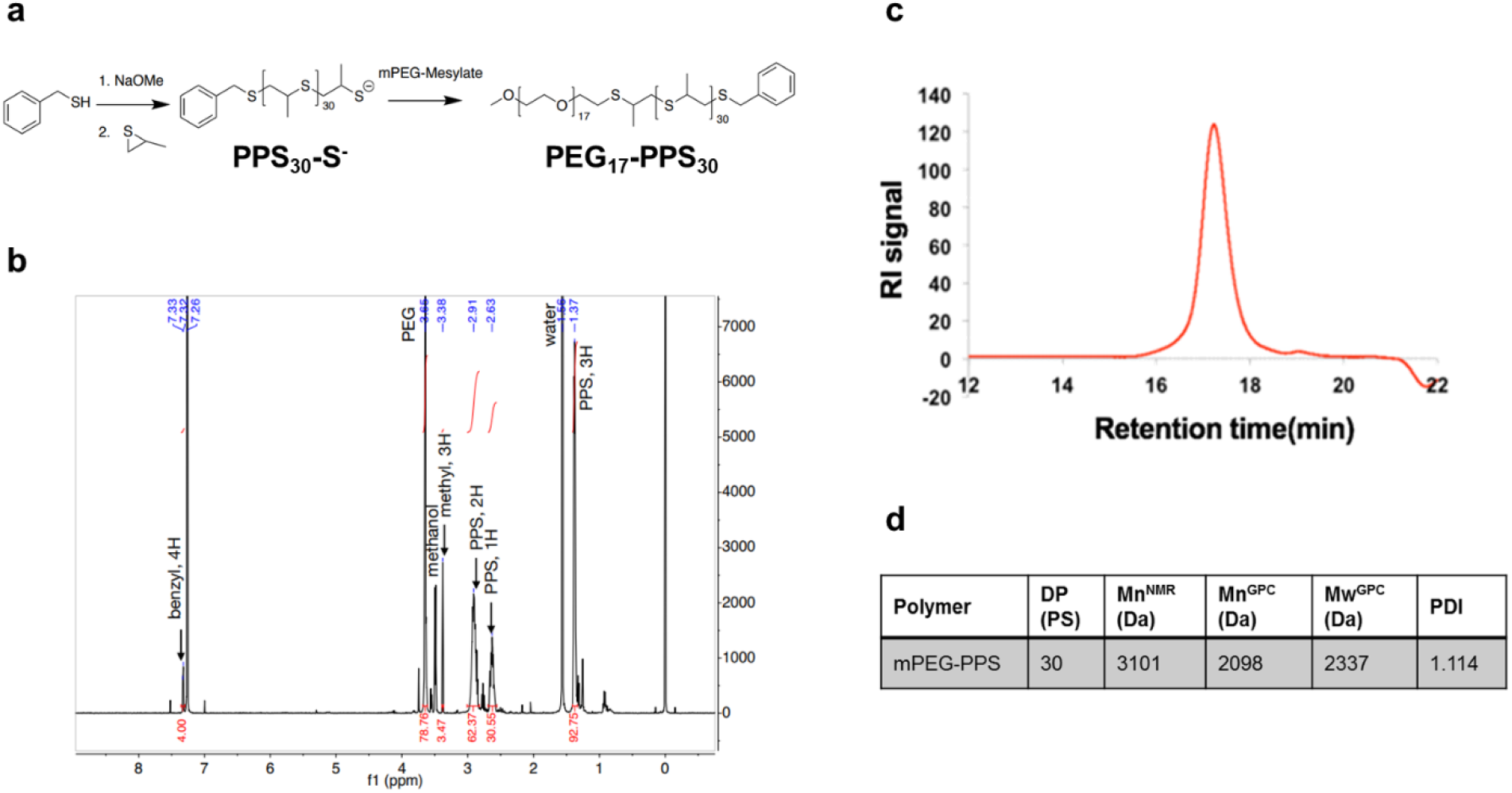
Synthesis and characterization of PEG-PPS. **a**, Synthetic route, **b**, ^1^H NMR spectrum, **c**, gel permeation chromatography (GPC) trace, and **d**, summary of physiochemical properties of PEG-PPS.

**Supplementary Figure S4.**
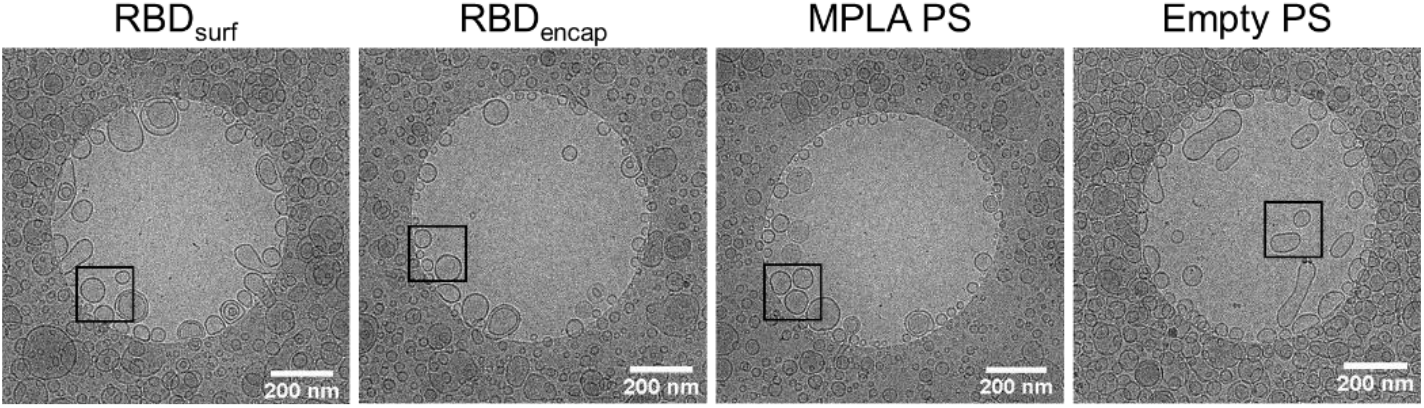
Additional cryoEM images of PS. Black box indicates magnified region in Figure 1b.

**Supplementary Figure S5.**
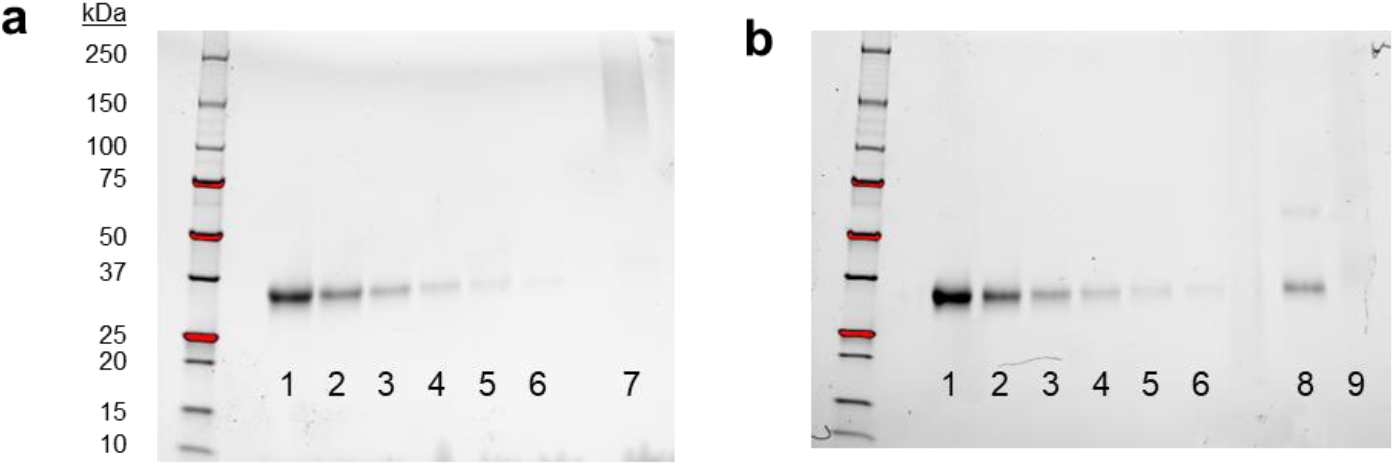
PS stability by SDS PAGE after > 180 d at 4 °C. SDS PAGE of **a**, RBD_surf_ and **b**, RBD_encap_. Lanes 1-6 represent RBD standard curve values of 400, 200, 100, 50, 25, and 12.5 µg/mL. Lane 7 contains of RBD_surf_ disrupted with Triton X. Lane 8 contains of RBD_encap_ disrupted with Triton X, and Lane 9 contains undisrupted RBD_encap_.

**Supplementary Figure S6.**
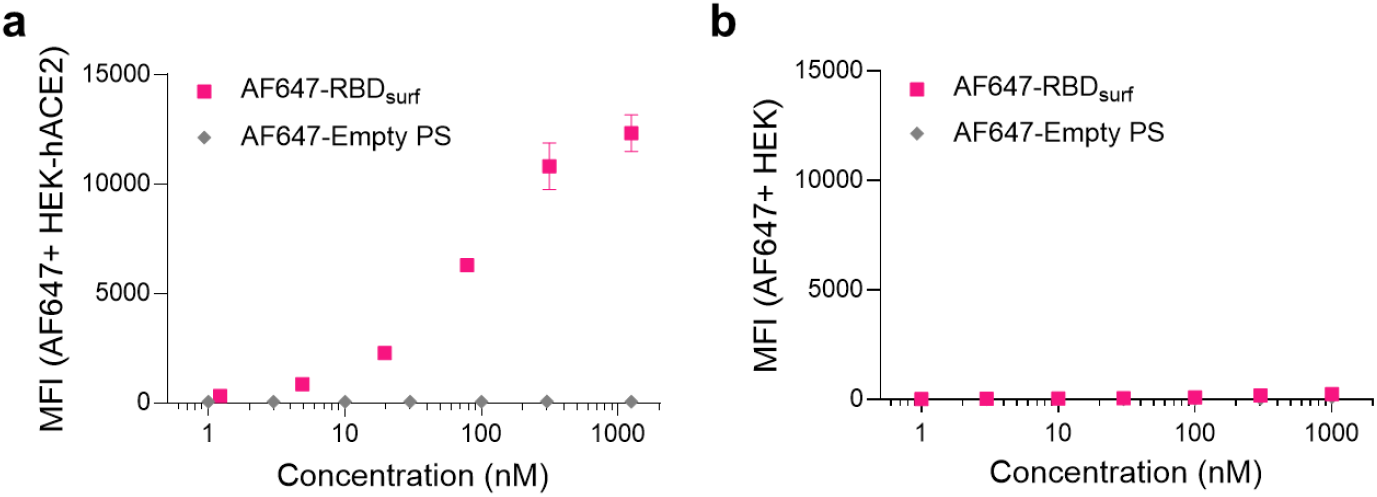
RBD binding to HEK-hACE2 and HEK-293 cells. **a**, Mean fluorescence intensity (MFI) of AF647-labeled RBD_surf_ and empty PS bound to HEK-hACE2 cells characterized by flow cytometry. **b**, MFI of AF647-labeled RBD_surf_ and empty PS indicating an absence of binding to control HEK-293 (HEK) cells. Data plotted as mean ± SD for n = 2 replicates.

**Supplementary Figure S7.**
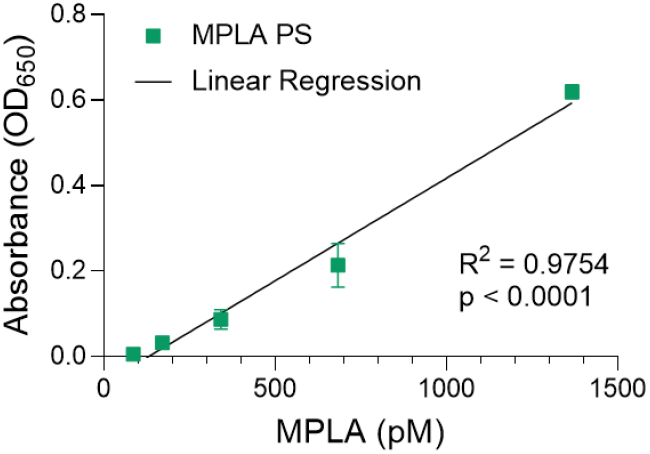
MPLA PS as a TLR4 agonist. Linear concentration-dependent stimulation of HEK-Blue™ TLR4 reporter cells with MPLA PS. Data plotted as mean ± SD for n = 3 replicates.

**Supplementary Figure S8.**
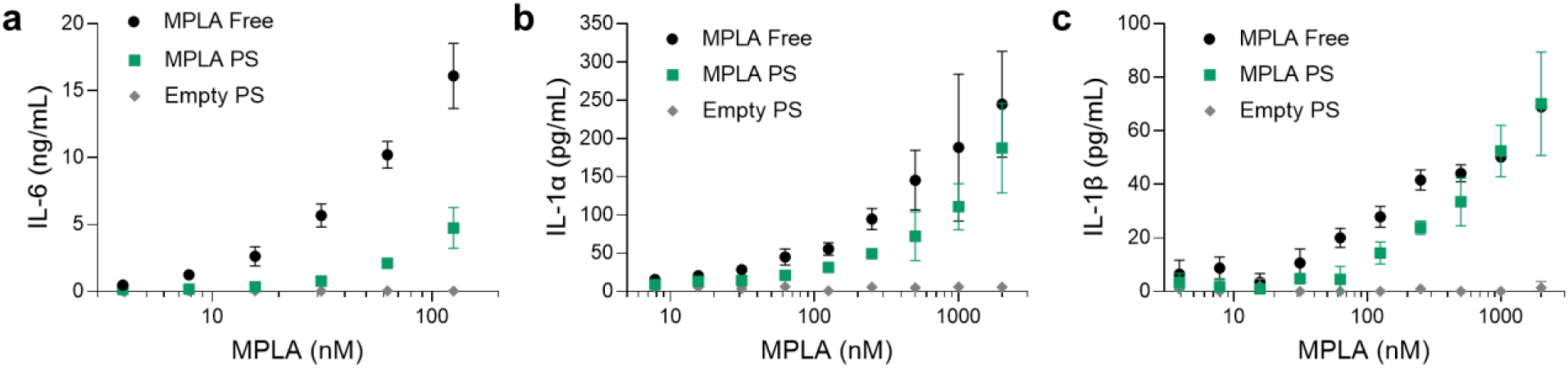
*In vitro* activity of MPLA PS. Dose-dependent secretion of **a**, IL-6, **b**, IL-1α, and **c**, IL-1β from cultured murine bone marrow-derived dendritic cells stimulated by free MPLA, MPLA PS, or empty PS. Data plotted as mean ± SD for n = 3 replicates.

**Supplementary Figure S9.**
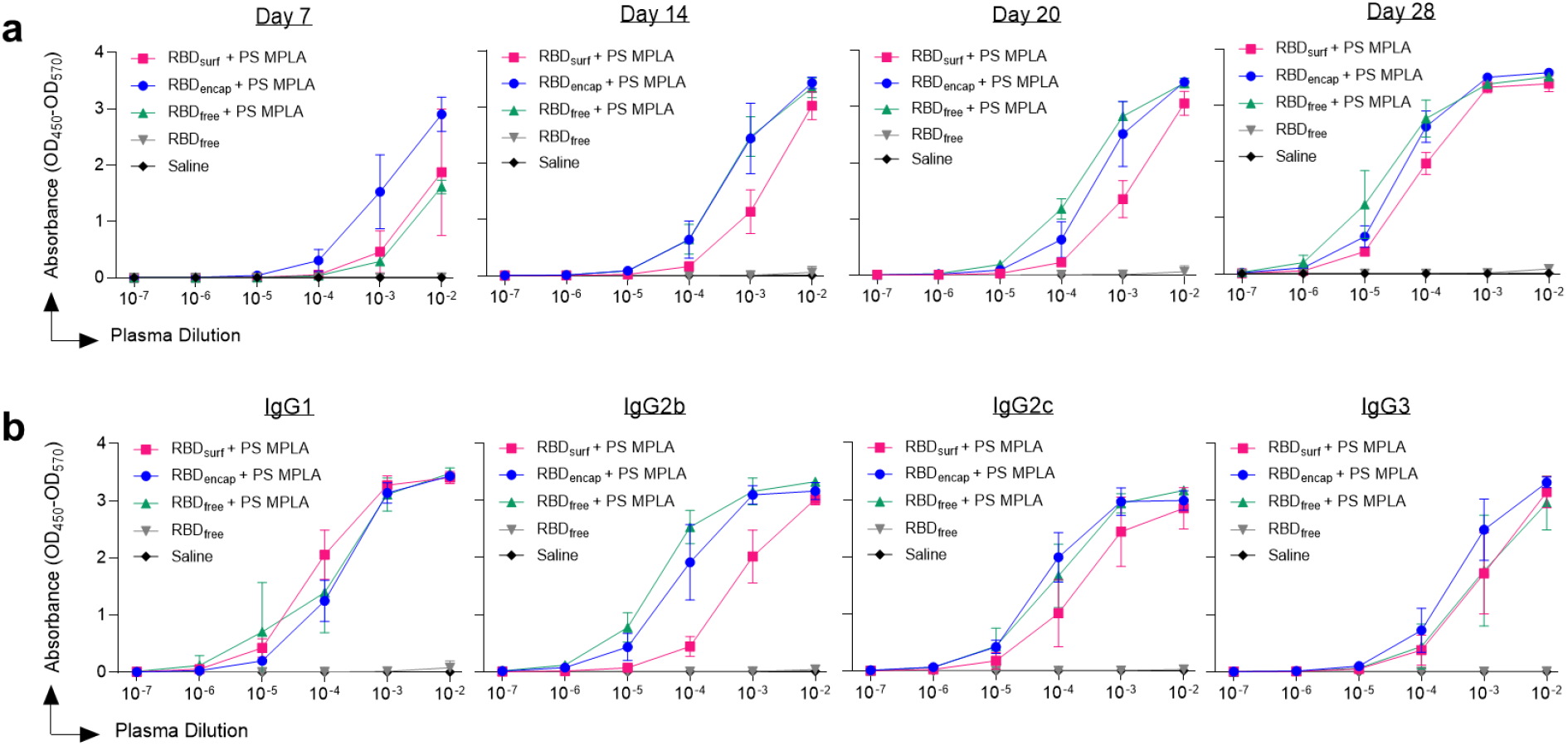
ELISA absorbance vs. dilution curves. Absorbance vs. dilution for RBD-specific ELISAs for **a**, total IgG over time and **b**, IgG subtypes on d28. Log-transformed curves were quantified by AUC in Figure 2. Data plotted as mean ± SD and represent 1 of 2 experiments with n = 5 mice each.

**Supplementary Figure S10.**
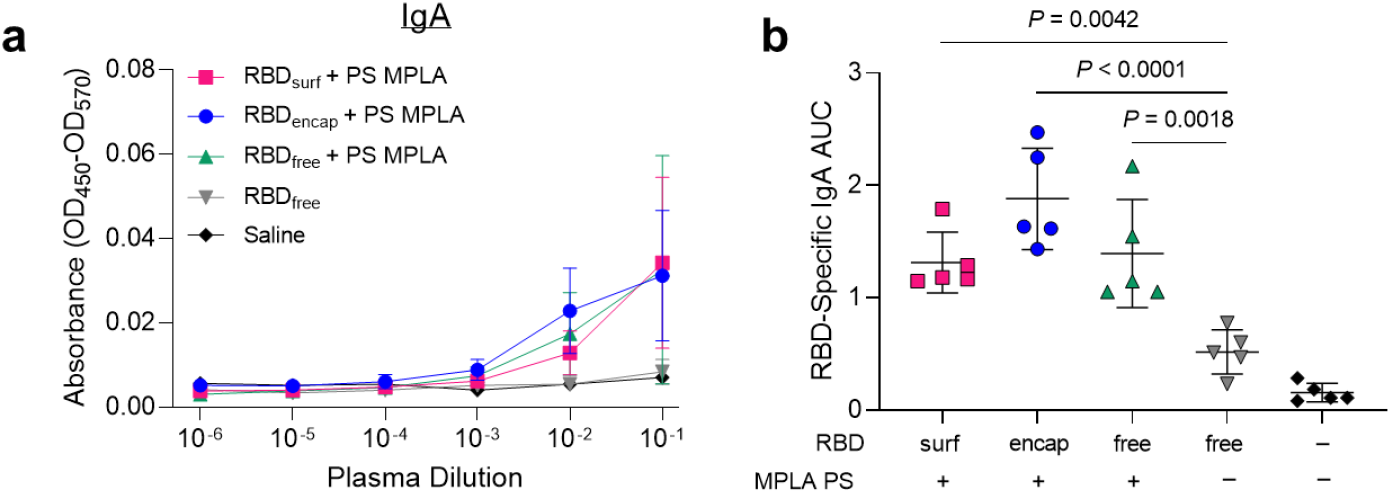
Presence of IgA antibodies. **a**, Absorbance vs. dilution for RBD-specific IgA ELISAs. **b**, AUC from (a). Data plotted as mean ± SD and represent 1 of 2 experiments with n = 5 mice each. Symbols in (b) represent individual mice. Comparisons to unadjuvanted RBD_free_ were made using one-way ANOVA with Dunn’s post-test.

**Supplementary Figure S11.**
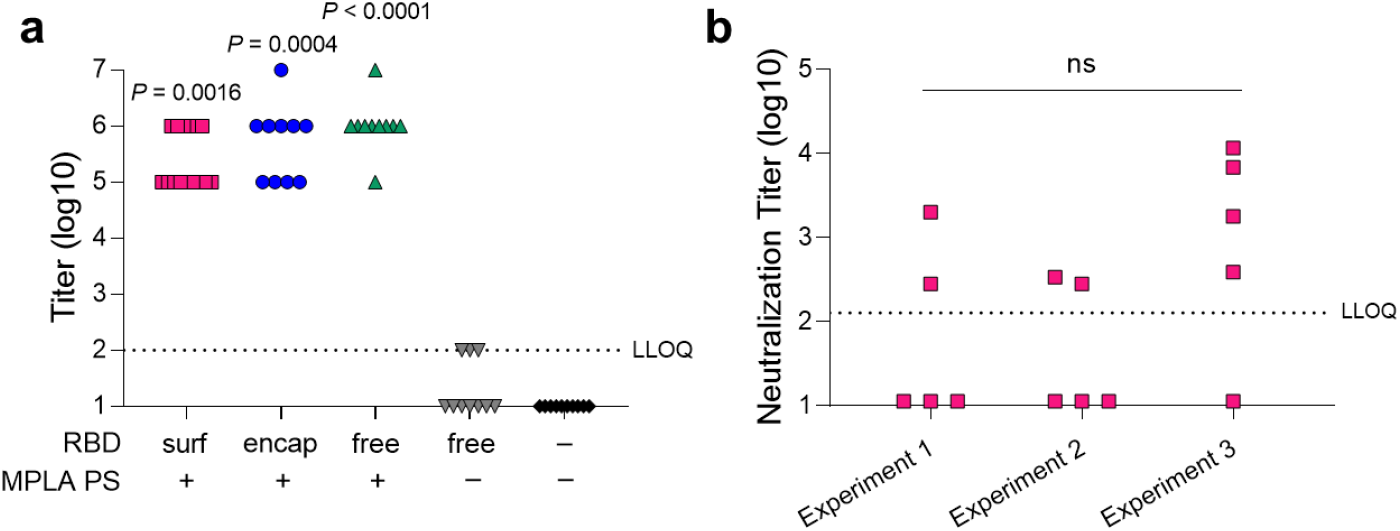
IgG antibody and viral neutralization titers. **a**, Aggregate RBD-specific IgG antibody titers 1 week post-boost based on ELISA. Values below the LLOQ (= 2) are plotted as LLOQ/2. Titers were determined as the -log of the lowest plasma dilution for which (OD450-OD570) – (average of blanks + 4 × standard deviation of blanks) > 0.01. *P* values represent comparisons to unadjuvanted RBD_free_. **b)** Viral neutralization titers for RBD_surf_ + MPLA PS across three different cohorts of n = 5 mice, indicating experiment reproducibility. Values below the LLOQ (= 2.11) are plotted as LLOQ/2.; ns p = 0.11. Symbols represent individual mice. Comparisons were made using a Kruskal-Wallis nonparametric test with Dunn’s post-test.

**Supplementary Figure S12.**
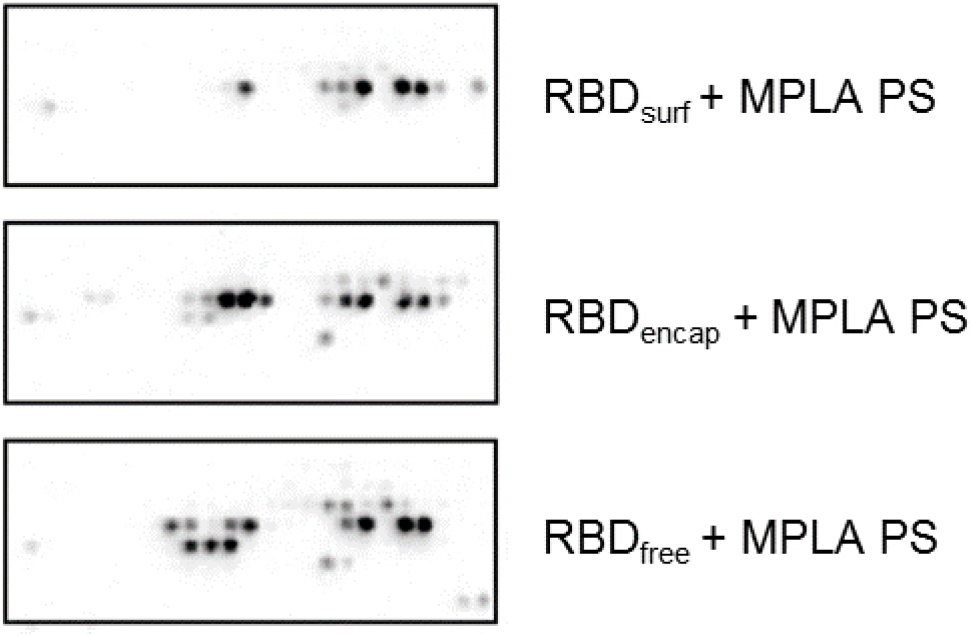
Representative peptide array images. Boxes represent region of peptide array specific to the RBD of the Spike protein. Peptide arrays quantified in Figure 3c.

**Supplementary Figure S13.**
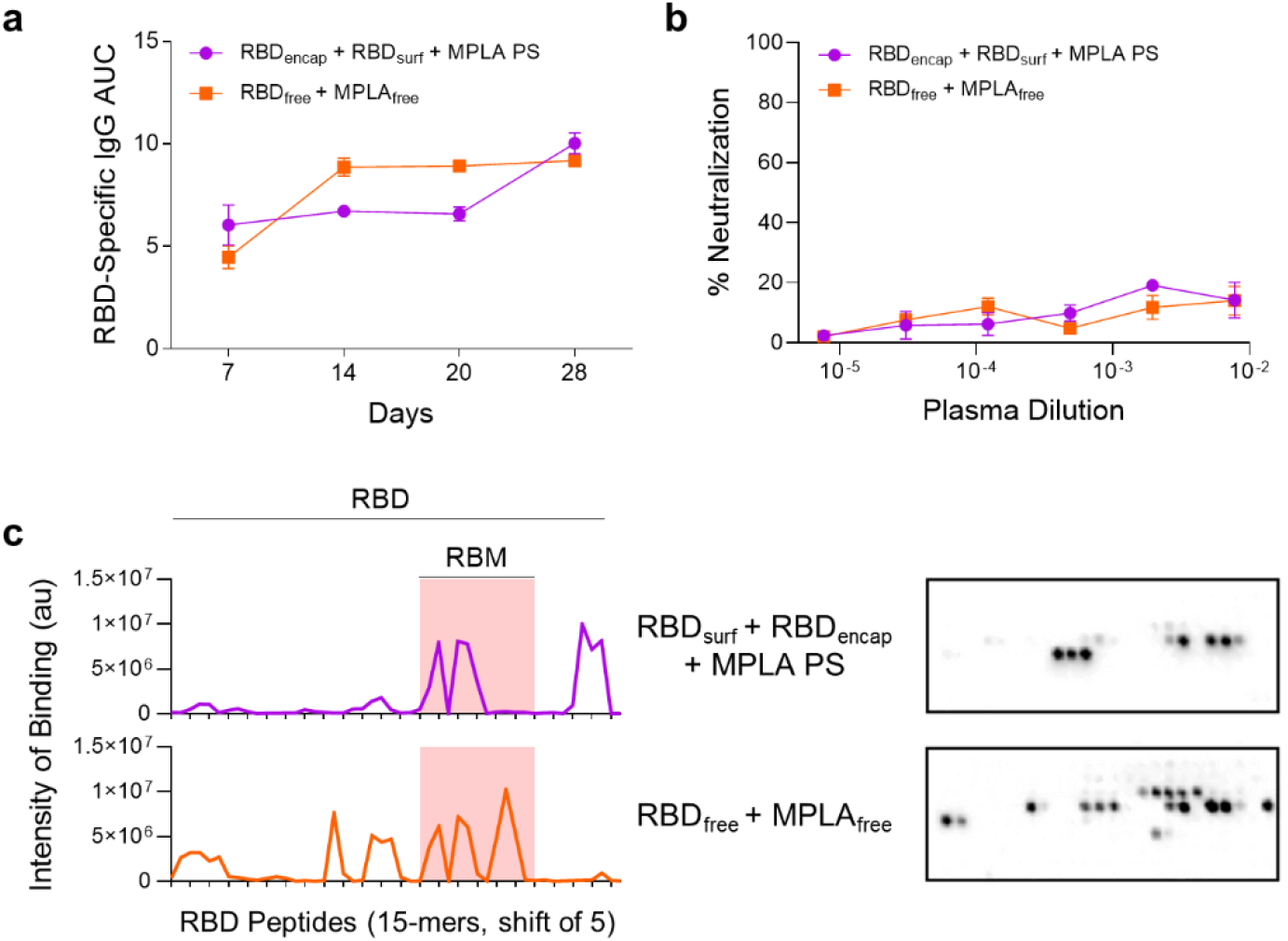
Analysis of plasma by mice vaccinated with RBD_surf_ + RBD_encap_ + MPLA PS and RBD_free_ + MPLA_free_. Mice received a priming dose on day 0 with a boost on day 21, and plasma was taken weekly to monitor production of RBD-specific antibodies. **a**, AUC of absorbance curve of RBD-specific IgG ELISAs for mice vaccinated with either 5 μg RBD_encap_ + 5 μg RBD_surf_ + MPLA PS or 10 μg RBD_free_ + MPLA_free_. Data plotted as mean ± SD for n = 5 mice. **b**, Neutralization of SARS-CoV-2 infection of Vero E6 cells *in vitro*. Data plotted as mean ± SEM for n = 5 mice. **c**, Epitope mapping using 15-amino-acid peptides with a 5-amino-acid shift, spanning the entire RBD sequence with representative images of peptide arrays.

**Supplementary Figure S14.**
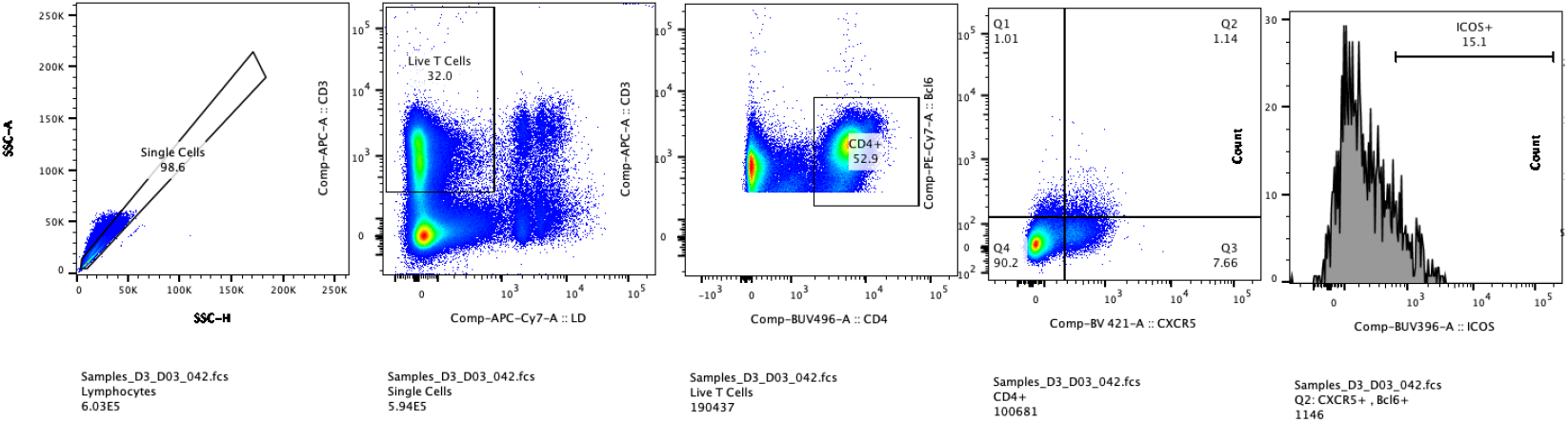
Representative T follicular helper cell gating strategy.

**Supplementary Figure S15.**
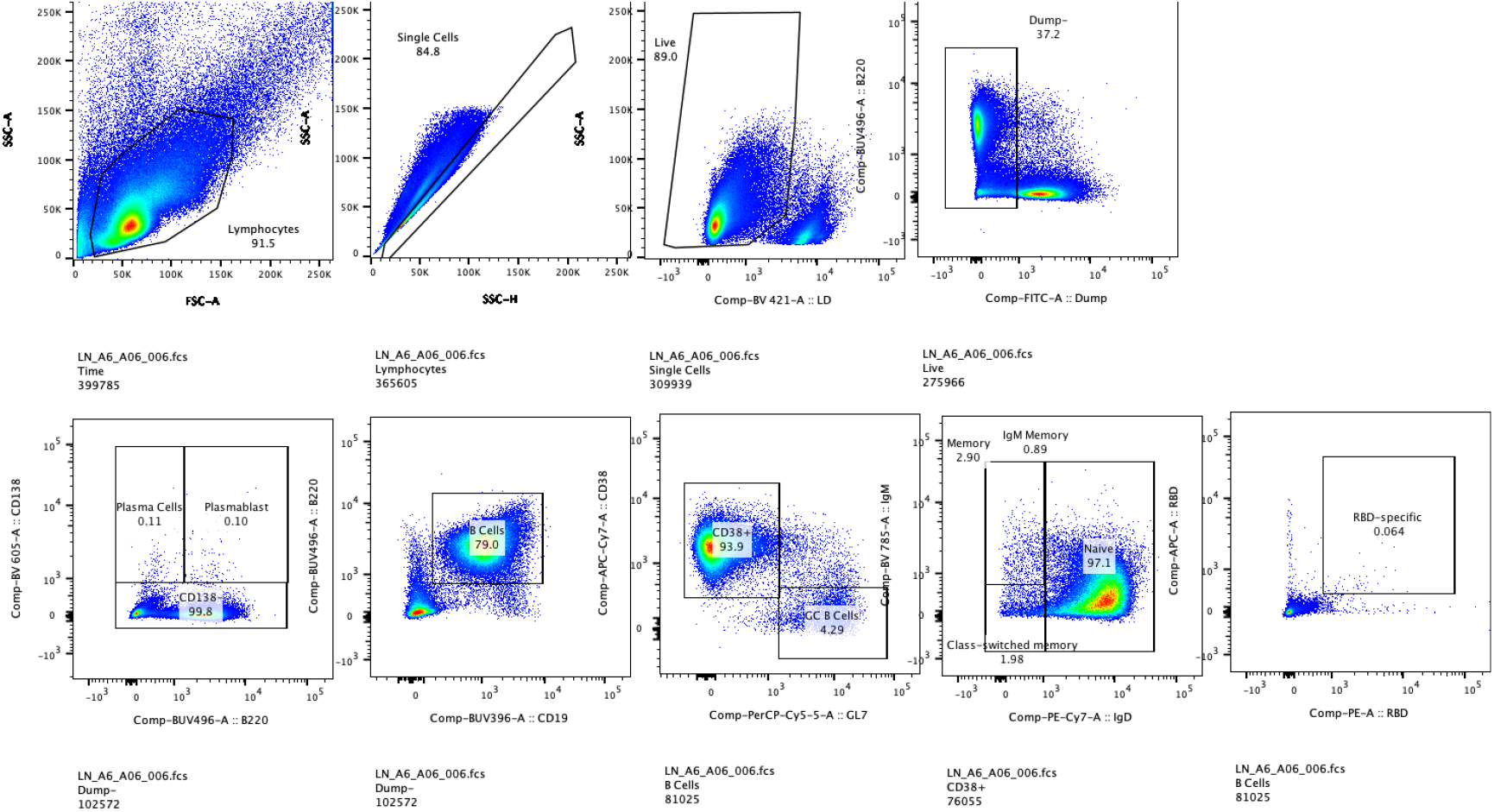
Representative B cell gating strategy.

**Supplementary Figure S16.**
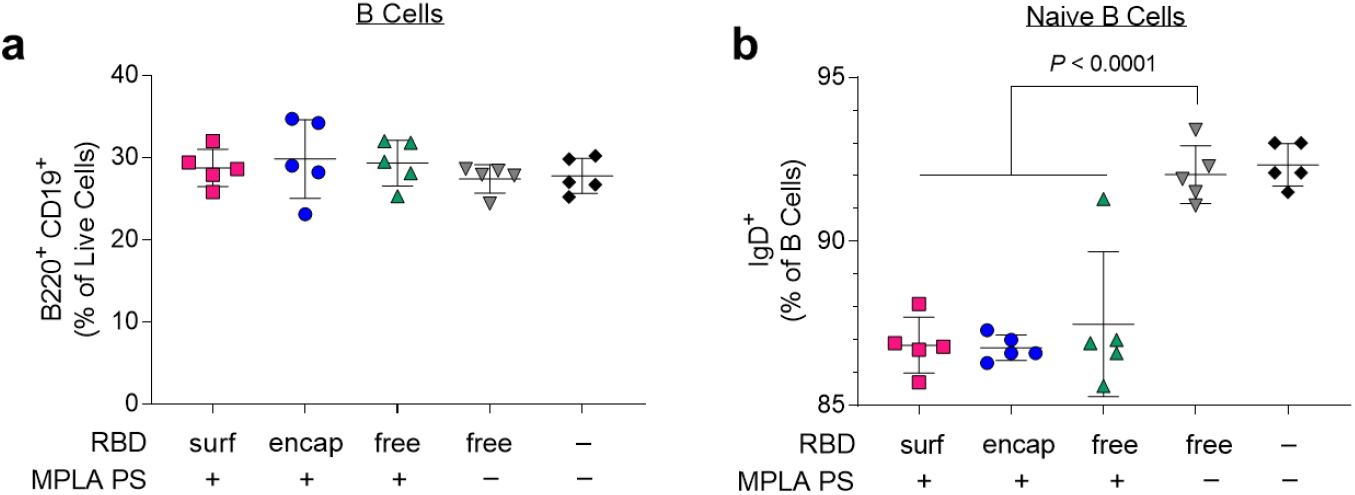
Total and naïve B cells in vaccinated mice 1 week post-boost. **a**, Total B cells (B220^+^ CD19^+^) and **b**, naïve B cells in dLNs. Data plotted as mean ± SD and represent 1 of 2 experiments with n = 5 mice each. Symbols represent individual mice. Comparisons to unadjuvanted RBD_free_ were made using one-way ANOVA with Dunn’s post-test.

**Supplementary Figure S17.**
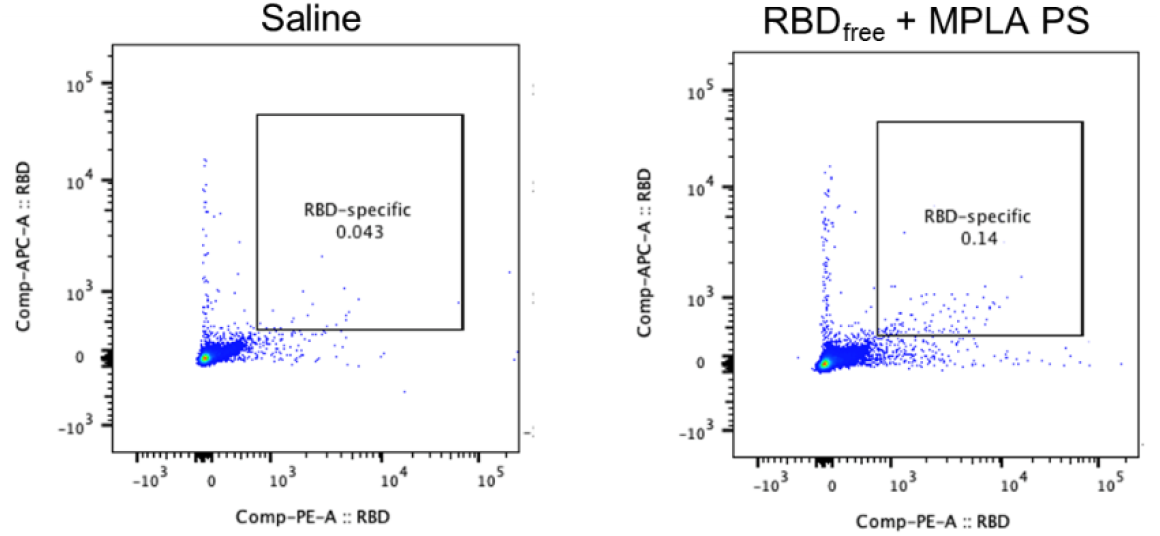
Representative tetramer staining.

**Supplementary Figure S18.**
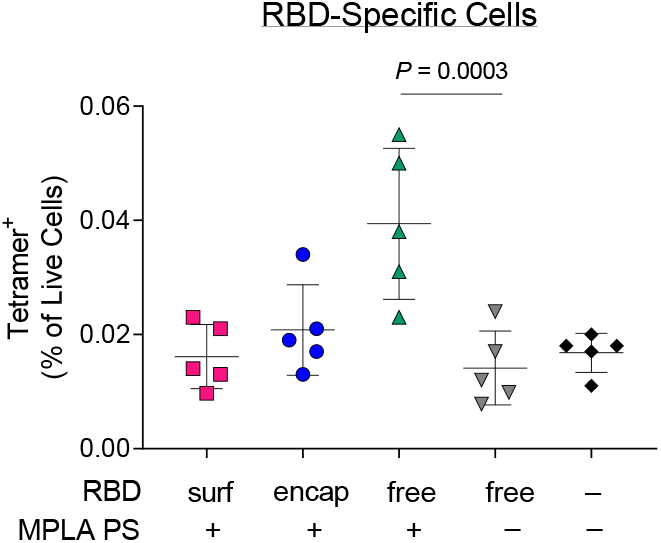
RBD-specific cells in vaccinated mice 1 week post-boost. Tetramer^+^ cells in dLNs on d28. Data plotted as mean ± SD and represent 1 of 2 experiments with n = 5 mice each. Symbols represent individual mice. Comparisons to unadjuvanted RBD_free_ were made using one-way ANOVA with Dunn’s post-test.

**Supplementary Figure S19.**
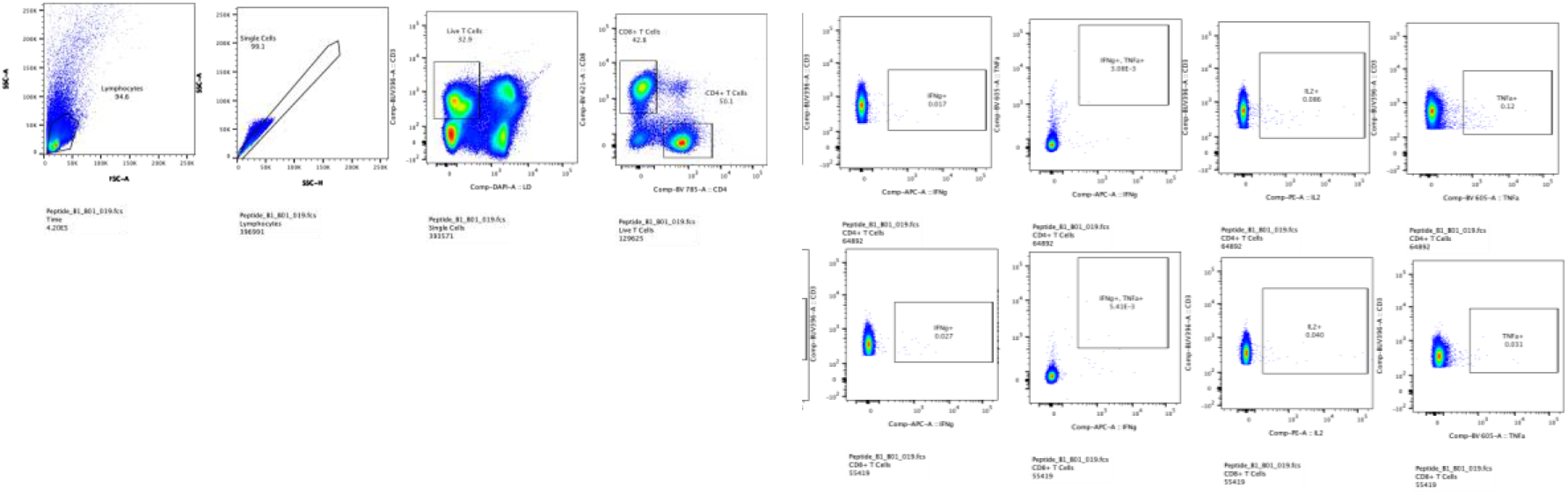
Representative intracellular cytokine gating strategy.

**Supplementary Figure S20.**
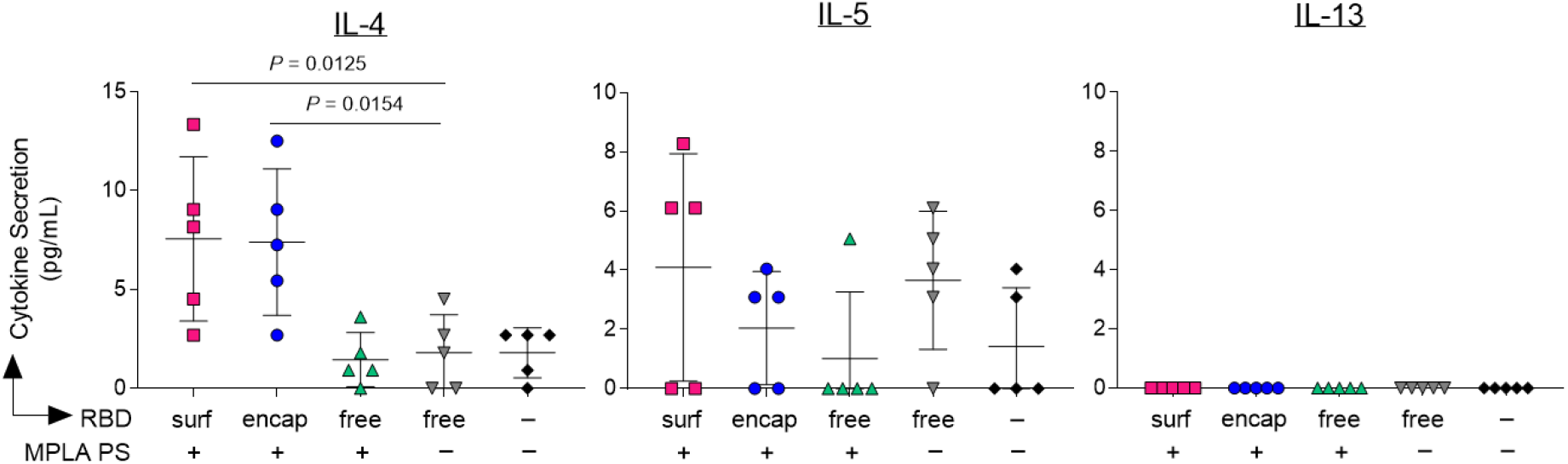
Th2-type cytokines secreted upon *ex vivo* stimulation with RBD. Lymph node cells isolated from the dLNs of PS vaccinated mice 1 week post-boost were restimulated *ex vivo* with full RBD protein. After 3 d, levels of IL-4, IL-5, and IL-13 secreted into the supernatant were measured. Data plotted as mean ± SD and represent 1 of 2 experiments with n = 5 mice each. Symbols represent individual mice. Comparisons to unadjuvanted RBD_free_ were made using one-way ANOVA with Dunn’s post-test.

**Supplementary Table S1.** Summary of loading capacities of PS.

|  | **Polymer (mg mL<sup>-1</sup>)** | **Cargo (mg mL<sup>-1</sup>)** | **Loading (wt%)** |
| --- | --- | --- | --- |
| RBD<sub>surf</sub> | 7.96 | 0.127 | 1.57 |
| RBD<sub>encap</sub> | 7.00 | 0.125 | 1.75 |
| MPLA PS | 3.49 | 0.241 | 6.46 |

Polymer concentration was determined by GPC, RBD concentration in RBD<sub>surf</sub> was determined by CBQCA protein quantitation assay, RBD concentration in RBD<sub>encap</sub> was determined by SDS PAGE, and MPLA concentration was determined by mass spectrometry.

**Supplementary Table S2.** Probes and antibodies for T_fh_ cell panel.

| **Marker** | **Color** | **Vendor** |
| --- | --- | --- |
| Viability Dye | eFluor 780 | Invitrogen 65-0865-14 |
| CD4 | BV496 | BD Horizon 612952 |
| CD3 | BUV737 | BD Optibuild 741788 |
| CD44 | PerCpCy5.5 | Invitrogen 45-0441-82 |
| CXCR5 | BV421 | Biolegend 145512 |
| ICOS | BUV396 | BD Horizon 565885 |
| Bcl6 | PE-Cy7 | Biolegend 358512 |

**Supplementary Table S3.** Probes and antibodies for RBD-specific B cell panel.

| **Marker** | **Color** | **Vendor** |
| --- | --- | --- |
| Viability Dye | Violet fluorescent reactive dye | Invitrogen L34964A |
| RBD-tetramer | PE | - |
| RBD-tetramer | APC | - |
| F4/80 (Dump) | FITC | Biolegend 123107 |
| CD11c (Dump) | FITC | Biolegend 117306 |
| Ly6c(Dump) | FITC | Invitrogen 53-5932-82 |
| Ly6g (Dump) | FITC | Invitrogen 11-9668-82 |
| CD4 (Dump) | FITC | Biolegend 100406 |
| CD8a (Dump) | FITC | Biolegend 100706 |
| B220 | BUV496 | BD Horizon 612950 |
| CD19 | BUV396 | BD Horizon 565965 |
| CD138 | BV605 | Biolegend 142531 |
| IgM | BV786 | BD Optibuild 743328 |
| IgD | PE-Cy7 | Biolegend 405720 |
| CD38 | APC-Cy7 | Biolegend 102727 |
| GL7 | PerCP-Cy5.5 | Invitrogen 46-5902-82 |

**Supplementary Table S4.** Probes and antibodies for restimulation panel.

| Marker | Color | Vendor |
| --- | --- | --- |
| Viability Dye | eFluor 455 (UV) | Invitrogen 65-0868-14 |
| CD3 | BUV395 | BD Horizon 563565 |
| CD4 | BV786 | BD Horizon 563331 |
| CD8 | BV421 | BD Horizon 563898 |
| IFNλ | APC | Biolegend 505810 |
| TNFα | BV605 | Biolegend 506329 |
| IL-2 | PE | BD Pharmigen 554428 |

## Chemical Synthesis and Characterization

### N_3_-PEG-PPS

N_3_-PEG-PPS was synthesized by first dissolving N_3_-PEG_24_-SH (1 eq, MW ∼1000 g mol^-1^; Nanosoft Polymers) in degassed, anhydrous THF and deprotonating the thiol group by addition of sodium methoxide (NaOMe; 1.1 eq) under nitrogen gas. Propylene sulfide (40 eq) was added by syringe, and the reaction proceeded until completion at the desirable degree of polymerization of PPS, as determined by ^1^H NMR. The polymer was precipitated multiple times in ice cold methanol to obtain the final product, N_3_-PEG_24_-PPS_40_, characterized by ^1^H NMR (400 MHz Bruker DRX spectrometer equipped with a BBO probe, using Topspin 1.3) and gel permeation chromatography (GPC; Tosoh EcoSEC size exclusion chromatography system with a Tosoh SuperAW3000 column). ^1^H-NMR (400 MHz, CDCl_3_) of N_3_-PEG-PPS, δ 1.37 (s, PPS, 3H), 2.63 (m, PPS, 1H), 2.91 (m, PPS, 2H), 3.39 (t, -CH_2_-N_3_, 2H), 3.65 (m, PEG).

### PEG-PPS

PEG-PPS was synthesized as previously described^1^. Briefly, benzyl mercaptan (1 eq.) in degassed, anhydrous THF (20 mM) was deprotonated with NaOMe (1.1 eq.). Under nitrogen protection, propylene sulfide (39 eq) was rapidly added by syringe, and the reaction proceeded until completion at the desirable degree of polymerization of PPS, as determined by ^1^H NMR. Subsequently, mPEG_17_-mesylate (synthesized in-house as previously described^1^) was added, and the mixture was allowed to react overnight. The polymer was precipitated multiple times in ice cold methanol to obtain the final product, PEG_17_-PPS_30_, characterized by ^1^H NMR and GPC. ^1^H-NMR (400 MHz, CDCl_3_) of mPEG-PPS, δ 1.37 (s, PPS, 3H), 2.63 (m, PPS, 1H), 2.91 (m, PPS, 2H), 3.38 (m, -OCH_3_, 3H), 3.65 (m, PEG), 7.32 (m, benzyl, 4H).

## Footnotes

1 Scott, E. A. et al. Dendritic cell activation and T cell priming with adjuvant- and antigen-loaded oxidation-sensitive polymersomes. Biomaterials 33, 6211-6219 (2012).

